# Deep learning to decode sites of RNA translation in normal and cancerous tissues

**DOI:** 10.1101/2024.03.21.586110

**Authors:** Jim Clauwaert, Zahra McVey, Ramneek Gupta, Ian Yannuzzi, Gerben Menschaert, John R. Prensner

## Abstract

The biological process of RNA translation is fundamental to cellular life and has wide-ranging implications for human disease. Yet, accurately delineating the variation in RNA translation represents a significant challenge. Here, we develop RiboTIE, a transformer model-based approach to map global RNA translation. We find that RiboTIE offers unparalleled precision and sensitivity for ribosome profiling data. Application of RiboTIE to normal brain and medulloblastoma cancer samples enables high-resolution insights into disease regulation of RNA translation.

## Main

RNA translation is an intricate process that involves the stepwise binding of the 40S and 60S ribosome subunits to RNA, along with multiple eukaryotic initiation factors and other cofactors.^1^ RNA translation is a major determinant of protein abundance and represents a core area of disease biology including cancer, where numerous genetic and non-genetic factors alter the composition of ribosomes, efficiency of translation, and fidelity of translation.^2^

To gain global insights into ribosome activity, ribosome profiling (Ribo-Seq) has become increasingly popular to determine the translational efficiency of mRNAs and detect non-canonical open reading frames (ORFs) and alternative proteoforms that have eluded standard analyses.^3,4^ The computational analysis of Ribo-Seq data has therefore become a cornerstone of research fields that rely on accurate ORF identification or RNA translation analyses, including *de novo* gene discovery, RNA regulation, proteogenomics, microprotein biology, and disease-focused research on therapeutic agents whose mechanism targets RNA translation. Yet, Ribo-Seq data analyses have been challenged by biases within the data that are caused by both biological (e.g., tissue type, cell lines vs. tissue samples) and technical (e.g., translation inhibitors, lab protocols) factors.^3^ Current computational approaches to Ribo-Seq have been developed on a central statistical test using manually curated features, where significant disagreement among tools reveals a lack of a “gold standard”.^5,6^

To address this problem, we have created RiboTIE, a transformer-based tool for the analysis of Ribo-Seq data. Our approach is based on current machine learning advances and enables the detection of translated ORFs within individual datasets without a pre-processing step, taking advantage of automated feature extraction to capture dataset-dependent correlations (**Fig. 1a**). RiboTIE processes ribosome information along the full transcript and can predict the presence of translation initiation sites (TISs) for each codon in any given RNA molecule. By evaluating every position as a candidate TIS, all possible ORFs on the transcriptome are scored. Importantly, RiboTIE only uses sequenced and mapped ribosome-protected fragments (RPFs) placed along the transcript and has no access to DNA sequence information (e.g., start codon) or ORF characteristics (e.g., length). Ribo-Seq data features reads of varying lengths, where the distribution of RPFs mapped on the reading frame of reference coding sequences (CDSs) versus out-of-frame is the default metric to assess data quality (**Extended Data Fig. 1**-2).^7^ Instead of adjusting or discarding any mapped reads based on this quality metric, RiboTIE processes all reads by position and length, which we determined improves its performance (**Extended Data Fig. 3**).

**Figure 1:**
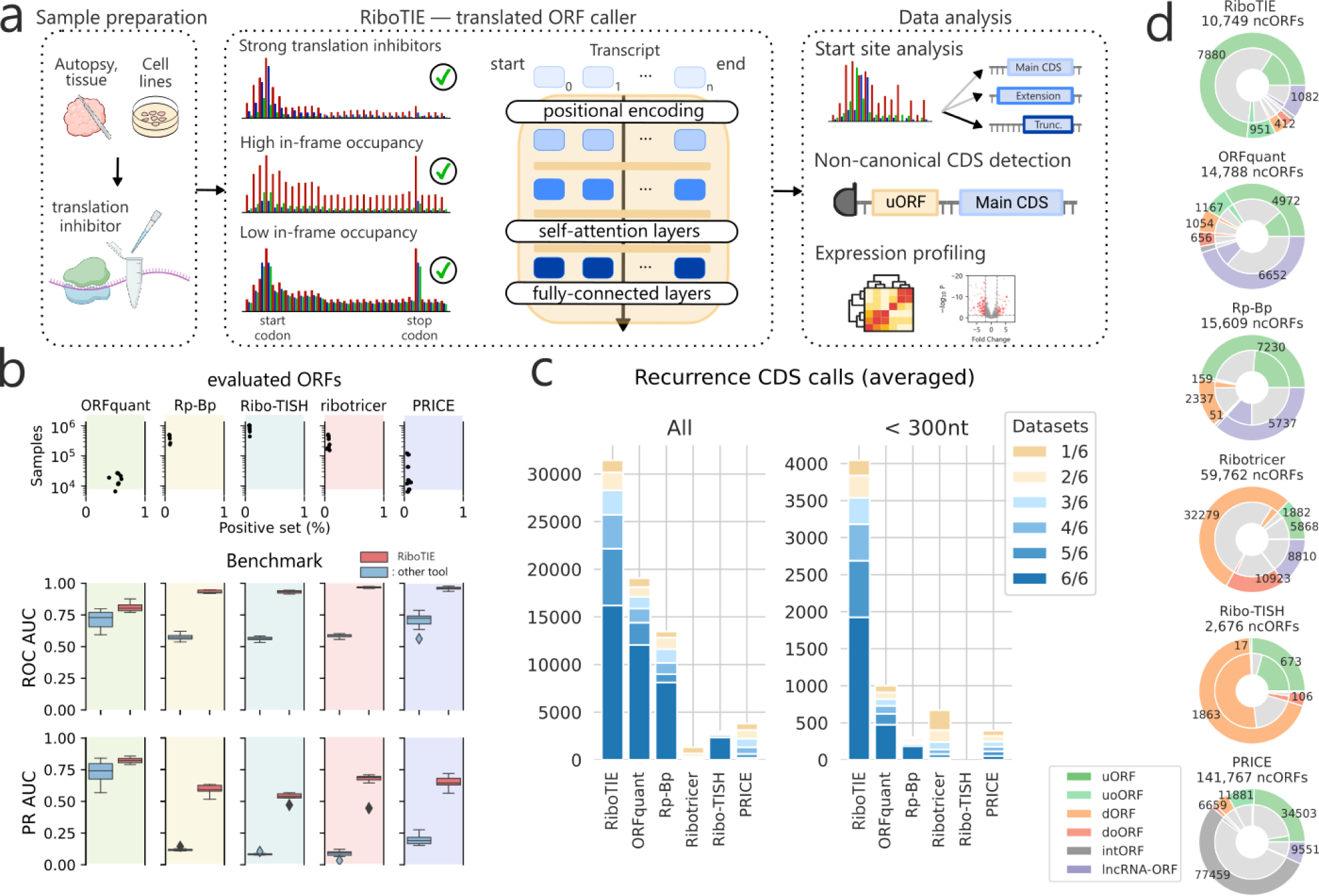
Machine learning to delineate RNA translation from Ribo-Seq data with RiboTIE. **a**, Schematic that outlines the flexibility and function of RiboTIE as a machine learning model (transformers) for ribosome profiling data. **b**, Benchmarking analyses featuring eight datasets. RiboTIE is compared with five other tools for translated ORF delineation from ribosome profiling. Precision recall (PR) or Receiver Operator Characteristic (ROC) Area Under the Curve (AUC) scores are compared on ORF libraries that are unique to each tool. **c**, A stacked barplot that reflects the number of called annotated CDSs (left, all; right, <300nt) by each tool for six replicate samples of pancreatic progenitor cells, the fraction of CDSs that are found in a certain number of replicates is represented as well. **d**, The total number of non-canonical ORFs (ncORFs) and each type of ncORFs called by each tool combining all predictions on the six replicate samples of pancreatic progenitor cells. The inner fractions represent ncORFs present in >4 datasets.

To assess the robustness of RiboTIE, we benchmarked our tool on eight Ribo-Seq experiments derived using varying treatment methods (**Extended Data Table 1**). Here, we compared RiboTIE against multiple common tools for translated ORF detection (ORFquant^8^, Rp-Bp^9^, Ribo-TISH^10^, Ribotricer^11^, and PRICE^12^), where scoring is performed on selection of ORFs evaluated by the other tools. We found RiboTIE to be more sensitive and precise on these ORF sets, as reflected by the Area Under the Receiver Operating Characteristics (ROC AUC) and Precision Recall (PR AUC) curve (F**ig. 1b**; **Extended Data Table 2**).

As the benchmark evaluation is potentially biased against RiboTIE due to the set of evaluated ORFs being curated by other tools, we compared the positive predictions of each tool in more detail by analyzing six biological replicates of pancreatic progenitor cells (**Extended Data Fig. 4**).^13^ We found that RiboTIE retrieved 64.9% more CDSs (31,431) as compared ORFquant (19,064), which obtains the second most calls for annotated CDSs (**Fig. 1c**). For smaller CDSs less than 300 bp in length, RiboTIE retrieves 300% more CDSs (4,043) as compared to ORFquant (1,001). RiboTIE reproducibly called 50% of annotated CDSs for each of the six datasets, where only an average of 4.2% of the calls are unique within a dataset (**Fig. 1c**).

We found that RiboTIE retrieved the largest quantity of annotated CDSs with non-canonical start sites across all six datasets (48; all CUG), where ORFquant (1; AUA) and PRICE (21; all CUG) are the only other tools to also feature non-canonical start codons. We also find notable differences between the number and types of non-canonical ORFs nominated by different tools (**Fig. 1d**). Notably, RiboTIE has a substantially higher fraction of upstream (overlapping) ORFs (u(o)ORFs) as compared to other tools. The fraction of internal ORFs (intORFs) and downstream (overlapping) ORFs (d(o)ORFs) is low with RiboTIE, in contrast to Ribo-TISH and Ribotricer, as prediction of these ORFs is known to be plagued by false-positive calls.^3,5^ Additionally, RiboTIE calls considerably fewer lncRNA-ORFs compared to other high-performing tools such as ORFquant and Rp-Bp. Nonetheless, for all three tools, the called lncRNA-ORFs follow similar distributions: ∼25% of called lncRNA-ORFs have TISs that overlap with protein-coding sequences, whereas ∼46% overlap with exons from protein-coding transcripts (**Extended Data Fig. 5**).

We next sought to apply RiboTIE to human tissue samples, where data quality may be more variable compared to cell line experiments. We evaluated 73 brain samples from both fetal (30) and adult (43) patients^14^ along with 15 medulloblastoma patient tissues^15^ (**Fig. 2a**). Notably, data quality is poor for some of the samples, with total in-frame read occupancies below 60% in 32 samples and below 50% in 12 samples (**Fig. 2a**). Across the 73 fetal and adult normal brain samples, RiboTIE made a total of 89,682 unique ORF calls (**Extended Data Fig. 4**), of which 39,828 (44.4%) were annotated CDSs and 16,253 ncORFs (18.1%) (**Fig. 2b**). This represented a substantial performance improvement relative to a much larger number of called ORFs (158,855, of which 28.9% are annotated CDSs and 30.8% ncORFs) previously reported for the same dataset through the RibORF software.^16^

**Figure 2:**
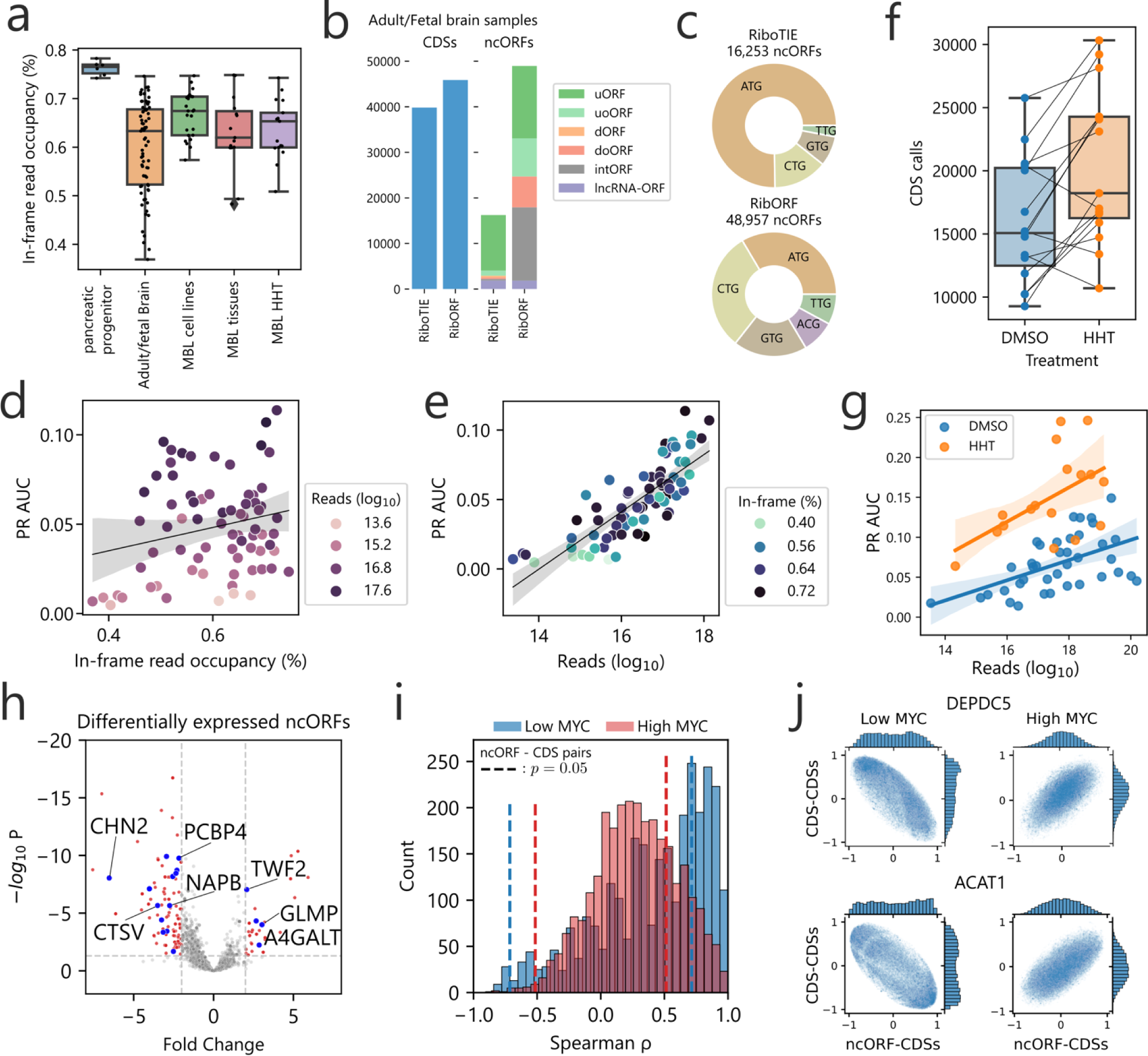
Application of RiboTIE to human normal tissues and brain cancer for improved analysis of RNA translation. **a**, Box plot showing the in-frame read occupancy (reads mapped to reading-frame vs. total reads within CDSs) for all data applied in this study (MBL: medulloblastoma). **b**, Bar plot displaying the combined number of unique calls for annotated CDSs and ncORFs on 73 adult/fetal brain samples as reported by the original paper^14^ (RibORF) and RiboTIE. **c**, A pie chart on the start codon distribution of all called ncORFs. **d**, Scater plot displaying the PR AUC performance of RiboTIE on adult/fetal brain samples as a function of mapped reads on the transcriptome and **e**, in-frame read occupancy. **f**, Number of CDSs called by RiboTIE outlined by both a scater plot and box plot for medulloblastoma cell lines treated with DMSO control or homoharringtonine (HHT). Identical cell lines are linked. **g**, Scater and fited linear regression plot on 30 DMSO (blue) and 15 HHT (orange) medulloblastoma samples. **h**, Volcano plot showing differential expression of called ncORFs of low MYC (n=8) as compared to high MYC (n=15) expressing medulloblastoma cell lines. Threshold lines denote p = 0.05 (y-axis) and |fold change| > 2 (x-axis). Blue dots accompanied by listed gene names are ncORFs confirmed by TIS Transformer. **i**, Histogram showing correlation existent between ncORFs and their matching CDSs for both low MYC (blue) and high MYC (red) cell lines. Threshold lines denote p = 0.05. **j**, Scater plots of Spearman rank correlations between the ncORF or downstream CDS and all other CDSs on the genome for both low and high MYC expression (SNAPC5/ACAT1).

Across these data, RiboTIE calls 36 CDSs with a non-AUG start codon as compared to 9 such instances by RibORF. For the ncORFs, the calls of RiboTIE were largely dominated by AUG start codons, whereas RibORF returns mostly non-canonical start codons (**Fig. 2c**). Interestingly, for the 73 evaluated brain samples that have varying in-frame read occupancies between 36% and 75% (**Fig. 2a**), we find only a slight and non-significant correlation with RiboTIE’s performance (spearman *ρ* = 0.178; *p* = 0.133) (**Fig. 2d**). As a strong correlation (Spearman *ρ* = 0.838; *p* = 2.5e − 20) does exist between the number of mapped reads within coding sequences and RiboTIE’s performance (**Fig. 2e**), our results indicate that reads that appear out-of-frame for technical and sample processing or quality reasons, are equally leveraged by RiboTIE to determine translated ORFs. In addition, there is an even stronger correlation when only considering the number of reads around the TISs (+/- 30nt) and RiboTIE’s performance (Spearman *ρ* = 0.947; *p* = 1.1e − 31).

Following this observation, we generated ribosome profiling data from medulloblastoma cell lines either treated with dimethyl sulfoxide (DMSO) or homoharringtonine (HHT). HHT blocks translation and concentrates RPFs around TISs. Compared to DMSO, we find an increased number of CDSs retrieved for 12 out of 15 pairs of HHT-treated cell lines (**Fig. 2f**). Including 15 medulloblastoma tissue samples from a previous study,^15^ we find HHT-treated cells to improve performance when incorporating the effect of read depth between samples (**Fig. 2g**).

To illustrate the role of ncORFs in medulloblastoma, we processed a total of 24 medulloblastoma cell line samples utilizing RiboTIE to evaluate differentially expressed ncORFs between samples with high (n=16) and low (n=8) MYC expression, which is used to classify distinct medulloblastoma subtypes.^17^ Across all datasets, a total of 3,638 ncORFs, of which 69.4% are upstream ORFs (uORFs), were selected for evaluation. We found 190 ncORFs with substantial alterations (|Fold Change| > 2; *p*_*adj*_< 0.05) in translational expression (**Fig. 2h**; **Extended Data Table 3**). We further integrated results from TIS Transformer,^18^ a machine learning tool previously developed by us to predict translated ORFs by RNA sequence context alone, to filter down candidate ncORFs by their sequence properties. Combined RiboTIE and TIS Transformer analyses revealed 22 candidate ncORFs with differential translation between medulloblastoma subtypes (**Fig. 2h**, annotated with blue dots; **Extended Data Fig. 6**). In the context of cancer biology, we identified that clustering of the ncORFs read counts correctly groups the medulloblastoma disease subtypes (**Extended Data Fig. 7**). However, predicted ncORFs are often positioned close to, or overlapping with, annotated CDSs, where the majority of ncORF-CDS pairs are positively correlated (**Fig. 2i**). We observed 38 ncORF-CDSs pairs that show a negative correlation (Spearman *ρ* < −0.71; *p* < 0.05) for low MYC expression and 25 pairs for high MYC expression (Spearman *ρ* < −0.50; *p* < 0.05) (**Extended Data Table 3**). Interestingly, we find 13 ncORF-CDS pairs that are inversely correlated between cell lines with low and high MYC expression (all *p* < 0.05), of which 3 are further selected by the TIS Transformer model (DEPDC5, ACAT1, MPHOSPH6). Further evaluating the SNAPC5 ncORF and CDS with all other CDSs of the genome highlights the difference in both distributions for low and high MYC (**Fig. 2j**).

In summary, we have developed RiboTIE as a best-in-class and highly versatile analysis tool approach that utilizes machine learning to process Ribo-Seq data. We have widely applied RiboTIE across 166 datasets with varying sequencing depths (2.7E+5 – 2.5E+8 reads; **Extended Data Table 1**), demonstrating its unique ability to handle data with low in-frame read occupancy and refine biological insights for disease subtypes of childhood medulloblastoma. RiboTIE is available as a Python package (see **Code Availability**) with pre-trained models that allow fast optimization times on new data. RiboTIE offers a new avenue to spearhead studies on translation start site analysis, non-canonical ORF detection and expression profiling of translated ORFs using Ribo-Seq data.

## Methods

### Medulloblastoma cell culture

All parental cell lines were obtained from the American Type Culture Collection (ATCC, Manassas, VA) or the Bandopadhayay lab (MB002, D425, D458). Cell lines were routinely verified via STR genotyping and tested for mycoplasma contamination using the Lonza MycoAlert assay (Lonza) CHLA-01-MEDR, Med2112 (expressing mCherry and luciferase), Med411 (expressing GFP and luciferase), and MB002 cells were maintained in Tumor Stem Media comprised of DMEM/F12 (1:1) with Neurobasal-A medium (Invitrogen) and supplemented with HEPES (1M, 0.1% final concentration), sodium pyruvate (1mM final concentration), MEM non-essential amino acids (0.1mM final concentration), GlutaMax (1x final concentration), B27 supplement (1x final concentration), human EGF (20ng/mL), human FGF-basic-152 (20ng/mL), and heparin solution (2ug/mL final concentration). D283, D341, D384, D425, D458, DAOY, R262, UW228 and CHLA-259 cells were maintained in DMEM supplemented with 10% FBS and 1% penicillin-streptomycin in a 5% CO2 cell culture incubator. ONS76 cells were maintained in RPMI 1640 supplemented with 10% FBS and 1% penicillin-streptomycin in a 5% C02 cell culture incubator.

### Medulloblastoma ribosome profiling data

Published medulloblastoma cell line (DMSO treated) and tissue data were obtained from the Short Read Archive (PRJNA957428, for cell lines) or dbGAP (phs003446, for tissues). Matched homoharringtonine ribosome profiling data was generated for D283, DAOY, ONS76, R262, UW228, D384, D458, D425, Med411, SUMB002, Med2112, CHLA-01-MEDR, and D341. Ribosome profiling was performed as detailed previously.^15^ In brief, cells were grown to 60-70% confluence. Cells were then pre-treated with 5ug/mL homoharringtonine for 10 minutes. Cells were then harvested, washed once in 1x cold PBS, and collected by centrifugation. Cells were lysed in a buffer of 20mM Tris HCl, 150mM NaCl, 5mM MgCl2, 1mM dithiotrietol, 0.05% NP-40 and 25U/mL Turbo-DNase I (Invitrogen). 2ug/mL of cyclohexamide was additionally added to the lysis buffer. Cleared lysates were quantified and 2.5U/ug of RNAse I was added for 45 minutes at room temperature for digestion. The reaction was quenched with an equal volume of 1U/uL Superase RNase Inhibitor (Ambion). Ribosome protected fragments (RPFs) were isolated using ultracentrifugation at 55,000RPM at 4C for 2 hours. RPF RNA was purified with a Zymo Direct-Zol kit (Zymo) and rRNA was depleted using the siTOOLS human riboPOOL kit (siTOOLS Biotech, Germany). Denatured RPFs were resolved on a 15% TBE-Urea gel (200V, 65 minutes) and the RPF band at 26-32 nt was excised and RNA extracted. After RNA precipitation, RNA was end-repaired with T4 PNK at 37C for 1 hour, cleaned up with RNA Clean and Concentrator kit (Zymo). The 3’ linker adapter was ligated (22C, 3hrs) using T4 RNA ligase I and R4 RNA ligase 2 deletion mutant. Linker reactions were removed with Rec J exonuclease (Lucigen) and 5’ deadenylase (NEB). cDNA was synthesized with EpiScript RT enzyme (Lucigen, 50C for 30 minutes). After clean-up with Exonuclease I and RNAse I and hybridase (Lucigen), cDNA was resolved on a 10% TBE-Urea gel (175V for 1 hour), and 75nt bands were excised and DNA was precipitated. Precipitated cDNA was circularized with CircLigase (Lucigen, 60C for 3hrs). Circular cDNA was amplified using 2x Phusion HiFi master mix (NEB) in a 11-14 cycle PCR reaction depending on cDNA yield. PCR products were precipitated with ethanol and 5M NaCl, and resolved on a 8% TBE gel (100V, 90 minutes). The 150nt band was excised, purified, quantified for DNA abundance using a DNA Qubit, and analyzed for library size using an Agilent TapeStation (Agilent). Libraries were then sequenced on an Illumina NovaSeq.

### Ribosome profiling pre-processing

All generated and publicly available (hESC cells: GSE144682; Medulloblastoma DMSO samples: PRJNA957428; Medulloblastoma HHT samples: PRJNA1077309) data were mapped and processed using the same pipeline. Ensembl assembly GRCh38 version 110 was used as the reference transcriptome. Trimming was performed using cutadapt version 4.4 filtering out reads smaller than 14nt (-m 14).^19^ Next, STAR version 2.7.11a^20^ was used to filter contaminant RNA and DNA using the special arguments “SeedSearchStartLmaxOverLread .5”, “outFilterMultimapNmax 1000”, “outFilterMismatchNmax 2”, and “outReadsUnmapped Fastx”, and to map the left out reads to the transcriptome using the flags “quantMode TranscriptomeSAM”, “outFilterMultimapNmax 10”, “outMultimapperOrder Random”, “outFilterMismatchNmax 2”, and “seedSearchStartLmaxOverLread 0.5”, and “alignEndsType EndToEnd”.^21^

For further processing with RiboTIE, the aligned read files are read where the number of mapped reads per read length and position on the transcripts is used to create the vector embeddings for the model (**Supplementary** Fig. 1**; Extended Data Fig. 3**). For ORFQuant, Rp-Bp, Ribo-TISH, Ribotricer and PRICE, the recommended default settings are applied (**Supplementary Methods**).

### Vector embeddings from mapped reads

RiboTIE processes mapped RPF counts along the full transcript in parallel. No pre-processing of other types of data, such as start codon or ORF information, are used to curate features or build a candidate ORF library. Instead, only the mapped position (5’-end) and length of mapped ribosome protected fragments is utilized (**Supplementary** Fig. 2-4). To allow computation with a transformer-based architecture, vector representations are calculated that represent this information for each position on the transcriptome. For a given position, the vector embedding **e_c_** is obtained from the read count *c* and a set of feed-forward layers *ϕ*. Existing tools use a similar methodology but apply various approaches to offset mapped reads as a function of their read length (**Supplementary Table 1-2**). Read counts are normalized across the transcript for numerical stability.

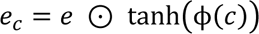

With 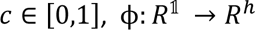, and *e* ∈ *R*^ℎ^. *h* is a hyperparameter of the model indicating the input dimension. In this paper, we explore the inclusion of ribosome read length information as part of the information applied to determine TIS locations. For a given transcript position, *e*_*l*_ is calculated using the read length fractions *l* between 20 and 41 following the equation:

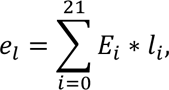

with ***E*** ∈ *R*^21×ℎ^ and **1** ∈ [0,1]^21^, where ∑ **1** = 1. The matrix ***E*** incorporates vector embeddings for read lengths 20–40 and is optimized as part of the training process. Note that ribosome data by read length is sparse and the majority of values in **1** are 0. After evaluation of different input embeddings (**Supplementary Table 3-4**), we find the optimal vector embedding for the model at each position to be *e*_**c**_ + *e*_*l*_.

### Model architecture and optimization

RiboTIE is created to map the translatome at single nucleotide resolution of samples using ribosome profiling data. To simplify the experimental learning objective of detecting translated ORF regions, we train a model to detect active translation initiation sites (TISs), denoting a binary classification problem. Open reading frames are afterwards derived using a greedy selection of the first in-frame stop codon.

Continuing upon our previous work on detecting TISs using transcript sequence information,^18^ we utilize an identical transformer framework for RiboTIE. RiboTIE makes use of the Performer architecture^22^ in order to allow calculation of the atention matrix spanning full transcript sequences. Evaluating various hyperparameters and training strategies, we obtained an optimal architecture featuring 212K weights (**Supplementary Methods, Supplementary Table 5-6, Supplementary** Figure 5-8). The model is optimized using a binary cross-entropy loss. To cover the full transcriptome without overfitting the machine learning model on the target labels, two models are trained on non-overlapping folds of the transcriptome. The training, validation and test sets are constructed from transcripts grouped per chromosome to ensure all transcript isoforms are present within the same set. Models are trained on the training set (chromosomes [3, 5, 7, 11, 13, 15, 19, 21, X], [2, 6, 8, 10, 14, 16, 18, 22, Y]) until a minimum loss on the validation ([1, 9, 17], [4, 12, 20] set is reached, where results are acquired from the test set ([2, 4, 6, 8, 10, 12, 14, 16, 18, 20, 22, Y, …], [1, 3, 5, 7, 9, 11, 13, 15, 17, 19, 21, X]). This approach, in which two models are optimized with the test sets covering the full transcriptome through non-overlapping folds is followed for all datasets. The outputs of both models are simply combined to atain predictions on all transcriptome positions.

### Data selection and result evaluation

A myriad of human ribosome profiling datasets were used covering a variety of tissues and treatment methods. Ensemble GRCh38 features a transcriptome totaling ∼250k transcripts and ∼431M transcript nucleotide positions, where annotated TISs (“start codon”) positions are utilized as the positive set when optimizing and evaluating the model.

When comparing RiboTIE with existing tools, performances are derived from all ORFs/TIS positions included by the ORF libraries constructed by such tools. This allows a one-on-one comparison as ORF libraries are unique to each tool. Otherwise, ROC/PR AUC values are obtained on all transcriptome positions (∼430M). As not all annotated TISs are expressed in each of the datasets, the maximum theoretical performance is affected.

The positive set of called ORFs is obtained from the full set of RiboTIE predictions by taking predictions above the threshold score of 0.15 and exclusion of ORFs with a start codon not equal to *TG (typically featuring ca. 10% of the complete set). For Ribotricer and RiboTISH, which feature large ORF libraries, the set of called translated ORFs is filtered down to be similar to the number of predictions made by RiboTIE. For Rp-Bp, the “--write-unfiltered” flag is used for benchmark results, to compare the output of the statistical method itself, whereas the filtered results are used for comparison of the positive sets on the human pancreatic cells. For the medulloblastoma data, ncORFs without in-frame stop codons were excluded from the set of ncORFs. For the results on the 73 adult/fetal brain samples, the set of called translated ORFs is filtered down to only include ORFs that have been called in more than one dataset to mirror the approach outlined by Duffy et al.^15^ with which a set of ORFs was derived using RibORF.

“lncRNA-ORFs” are defined for ORFs on transcripts with no annotated CDSs that furthermore do not share their genomic start or stop site with any annotated CDSs. ORFs existent on transcripts that share their genomic start or stop site with an annotated CDS are classified as “CDS variant”.

### Mapping differential expression and correlations between ncORF-CDS pairs

From a total of 26,437 ncORFs predicted by RiboTIE on all 24 medulloblastoma datasets, including u(o)ORFs, d(o)ORFs, intORFs and lncRNA-ORFs, a subset of 5,436 ORFs is obtained by selecting predictions that are present in at least 5 experiments. Combining this set with all canonical CDSs, differential expression is performed using the default workflow offered by PyDESeq2 0.4.4, where the number of mapped reads between samples that feature low (8) and high (15) expression of the MYC gene are compared. Differentially translated ncORFs were selected considering significance (*p*_*adj*_< 0.05) and the degree of change (|Fold Change| > 2). A further filtered down list of ncORFs is given by applying the predictions from TIS transformer as an additional condition (model output > 0.01).

Correlations were calculated on the set of 5,436 ncORFs and their CDS counterparts. Here, the Spearman ρ coefficients and p-values are calculated from mapped ribosome reads using the Transcripts Per Million (TPM) normalization method. A further filtered down list of anti-correlated ncORFs-CDS pairs is derived using TIS transformer (model output > 0.01).

### Data availability

The datasets generated during the current study are available in the Gene Expression Omnibus repository under accessions PRJNA1077309 (Medulloblastoma HHT/DMSO samples). Other analyzed datasets involve those on the Gene Expression Omnibus under accession PRJNA604580 (pancreatic progenitor cells) and on dbGAP under accession phs003446 (Medulloblastoma tissue samples) and phs002489 (adult/fetal brain samples).

## Code availability

RiboTIE (DOI: 10.5281/zenodo.10689717) is implemented in Python and is available through GitHub (https://github.com/jdcla/RIBO_former; future location https://github.com/jdcla/RiboTIE) and PyPI (https://pypi.org/project/transcript-transformer/).

## Acknowledgements

We thank Pratiti Bandopadhayay and Joelle Straehla for sharing medulloblastoma cell lines. We thank Sebastiaan van Heesch for helpful comments on the manuscript. J.R.P. acknowledges funding from the National Institutes of Health/National Cancer Institute (K08-CA263552-01A1), the Alex’s Lemonade Stand Foundation Young Investigator Award (#21-23983), the St. Baldrick’s Foundation Scholar Award (#931638), the DIPG/DMG Research Funding Alliance, the Book for Hope Foundation, the Yuvaan Tiwari Foundation, the Hyundai Hope on Wheels Foundation, the ChadTough Defeat DIPG Foundation, the Andrew McDonough B+ Foundation, the Curing Kids Cancer Foundation, and a Collaborative Pediatric Cancer Research Awards Program/Kids Join the Fight award (#22FN23). G.M. acknowledges support from Novo Nordisk.

## Author Contributions

Conceptualization: J.C., J.R.P., G.M.; methodology, J.C., J.R.P., I.Y., G.M.; formal analysis, J.C.; investigation, J.C., J.R.P., I.Y., G.M.; resources, J.R.P., G.M.; data curation, J.R.P., J.C.; writing - original draft, J.C., J.R.P., G.M.; writing - review & editing, J.C. and J.R.P. with input from all authors; visualization, J.C., J.R.P., G.M.; supervision, J.R.P., G.M.; project administration: J.C., J.R.P., G.M..; funding acquisition, J.R.P., G.M.

## Extended Data Tables

Extended Data Table 1: Datasets used in this study

Extended Data Table 2: Comparative performances of RiboTIE with different tools

Extended Data Table 3: Differentially expressed ncORFs between medulloblastoma cell lines with high and low MYC expression.

Extended Data Table 4: Read counts for CDSs across medulloblastoma cell lines

## Extended Data Figures

**Extended Data Figure 1:**
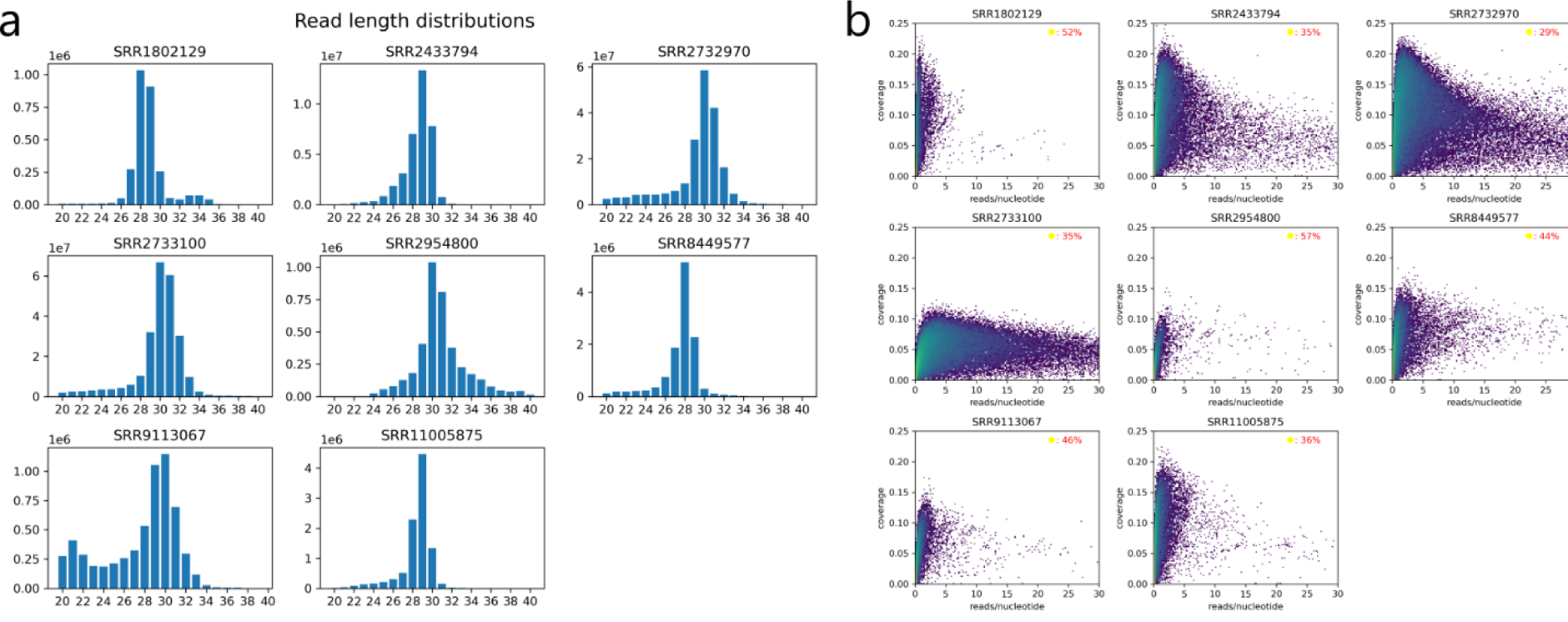
Benchmark datasets characteristics. **a**, Read length distributions for the benchmark datasets for reads mapped to the genome. The most abundant read length is generally around 29 nucleotides. **b**, 2D histogram of transcript-based coverage (y-axis) and reads mapped (x-axis). The number of mapped reads are normalized by transcript length (reads/nucleotide). The coverage is calculated based on the percentage of the transcript positions that have at least one read mapped by their 5’ position. The color map follows a logarithmic scale and is identical for all datasets. For each of the benchmark datasets, the percentage of transcripts in the transcriptome with no reads mapped is given (top-right corner).

**Extended Data Figure 2:**
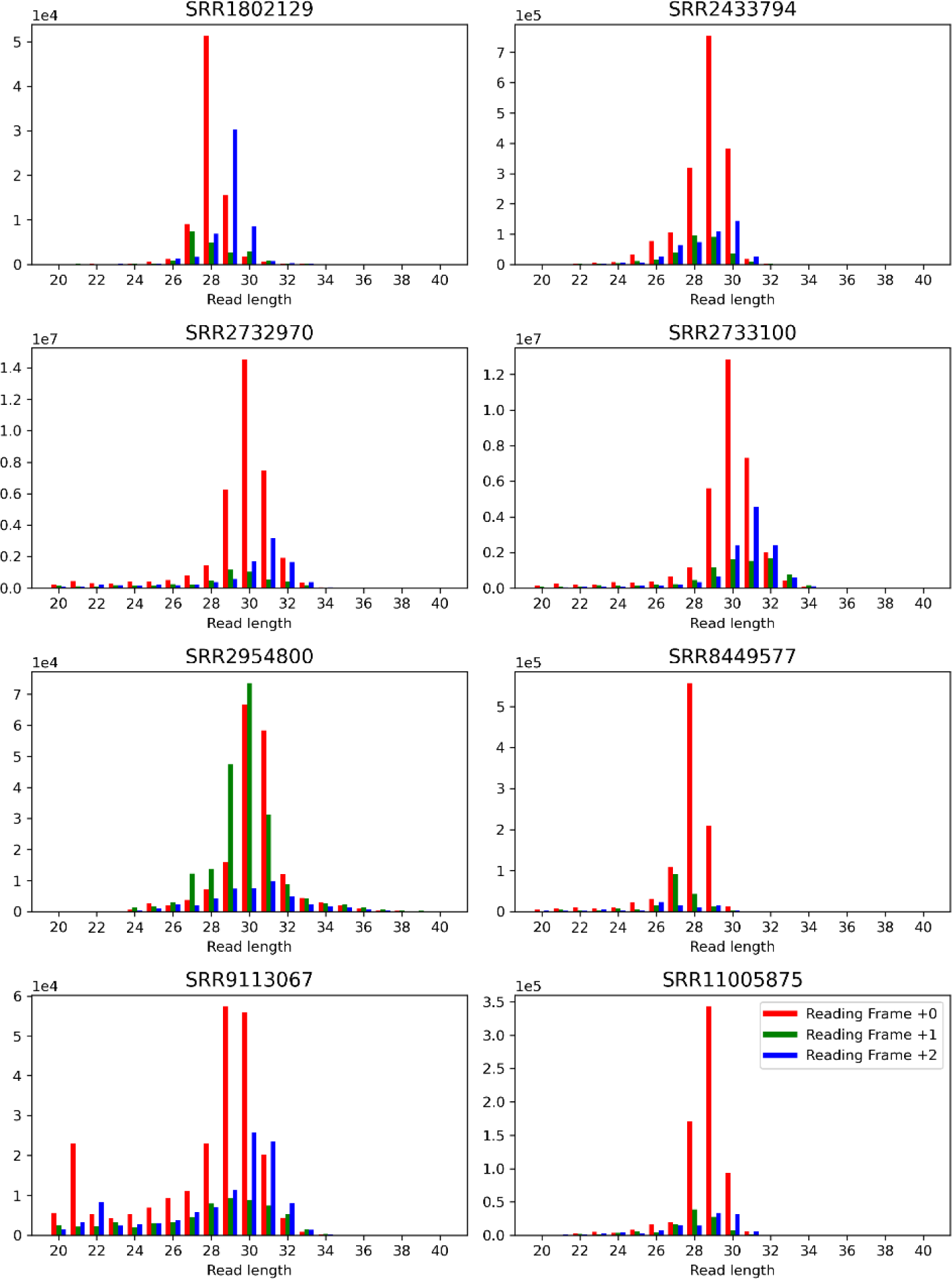
Read length counts binned by reading frame offset for all benchmark datasets. Reads are mapped by their 5’ positions. The figure highlights the skewed abundance of reads as influenced by the reading frame of the neighboring translation initiation site. Similar plots have been used to filter or offset the mapping position of reads in relation to their length. Read counts are taken by only evaluating translation initiation sites of coding sequences within the consensus coding sequence (CCDS) library. A window of 20 nucleotides upstream and 40 nucleotides downstream is taken to calculate the total read counts.

**Extended Data Figure 3:**
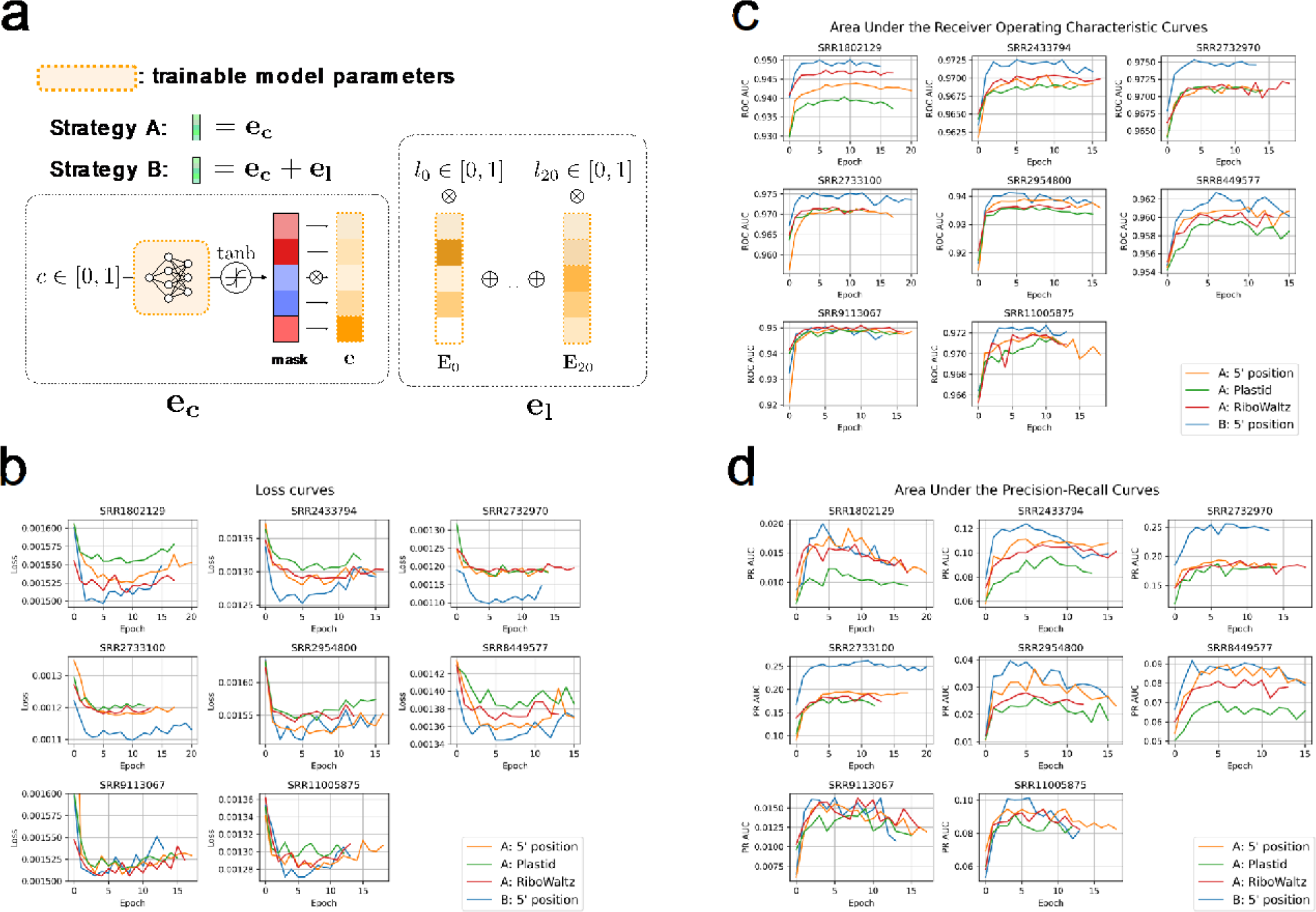
RiboTIE performances for different input token strategies and datasets. **a**, Illustration of two different strategies for constructing the RiboTIE input vector. Strategy A: the normalized read count is fed into short feed-forward neural network, where the output is used for an element-wise multiplication with a single vector embedding. Strategy B: vector embeddings are optimized for each read length. For a given input, read length embeddings are multiplied by the fractional representation of that read length at that position. Strategy B takes the sum of input vectors derived from the read count and read lengths. (**b**, **c, d**), Scores are calculated on the test set after selection of the model with the minimum validation loss. For each dataset and strategy, the cross-entropy loss, area under the receiver operating characteristic curve (ROC AUC), and area under the precision-recall curve (PR AUC) are given. Results indicate the relevance of read length information for the prediction of translation initiation sites using ribosome profiling data, especially for datasets featuring a higher read depth. All strategies are evaluated using the same model architecture and training/validation data. Strategy A has been evaluated for reads mapped by their 5’-end and reads offset based on read length information utilizing two different tools (Plastid, RiboWaltz).

**Extended Data Figure 4:**
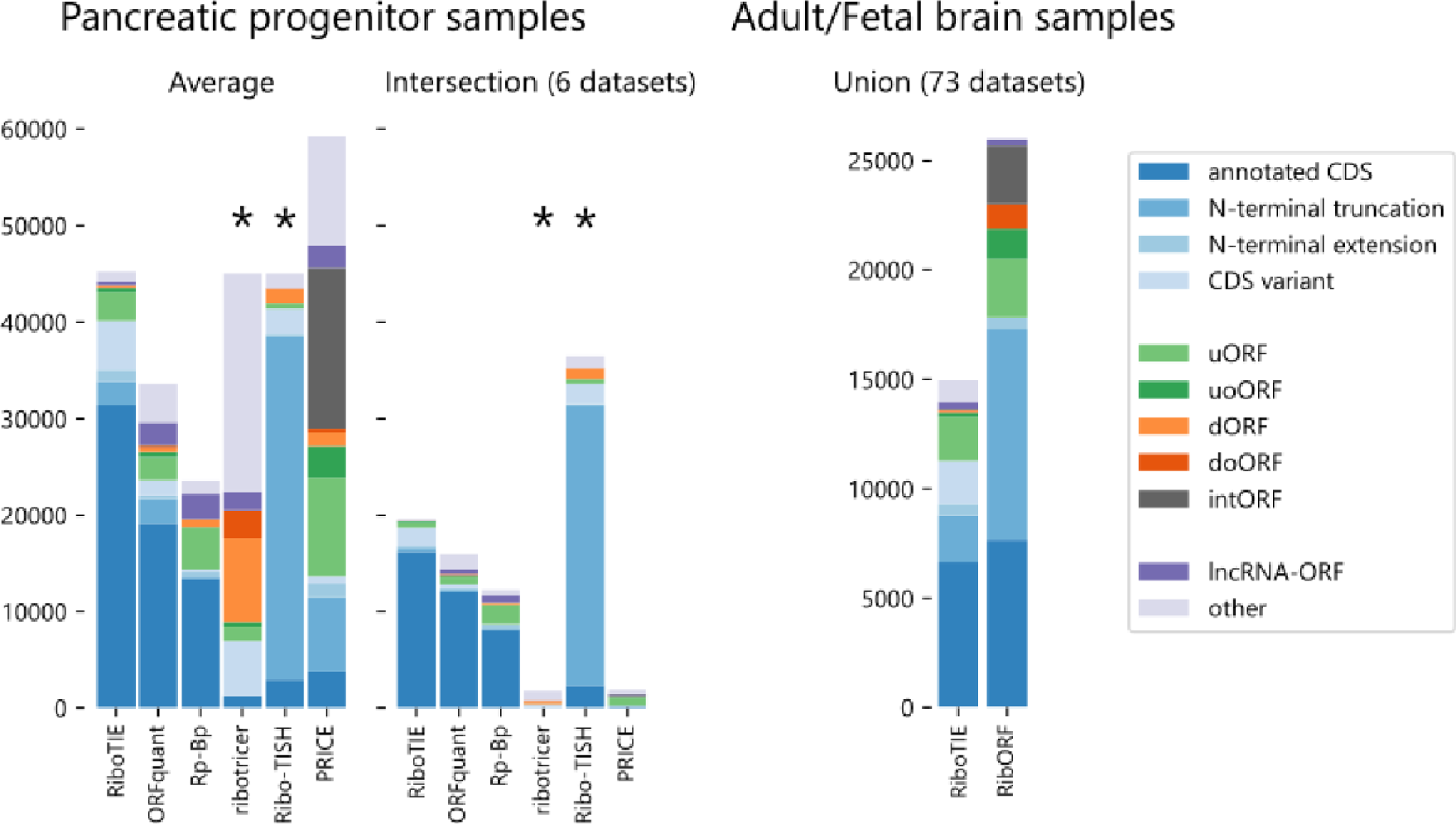
Stacked bar plot denoting the number of ORFs for each type within the positive set of various tools on the pancreatic progenitor cells and adult/fetal brain samples. Tools tagged with “*” (ribotricer/Ribo-TISH) give output predictions on all ORFs within their ORF libraries. As such, a positive set with an identical size to that of RiboTIE was selected for comparison by taking the top scoring predictions.

**Extended Data Figure 5:**
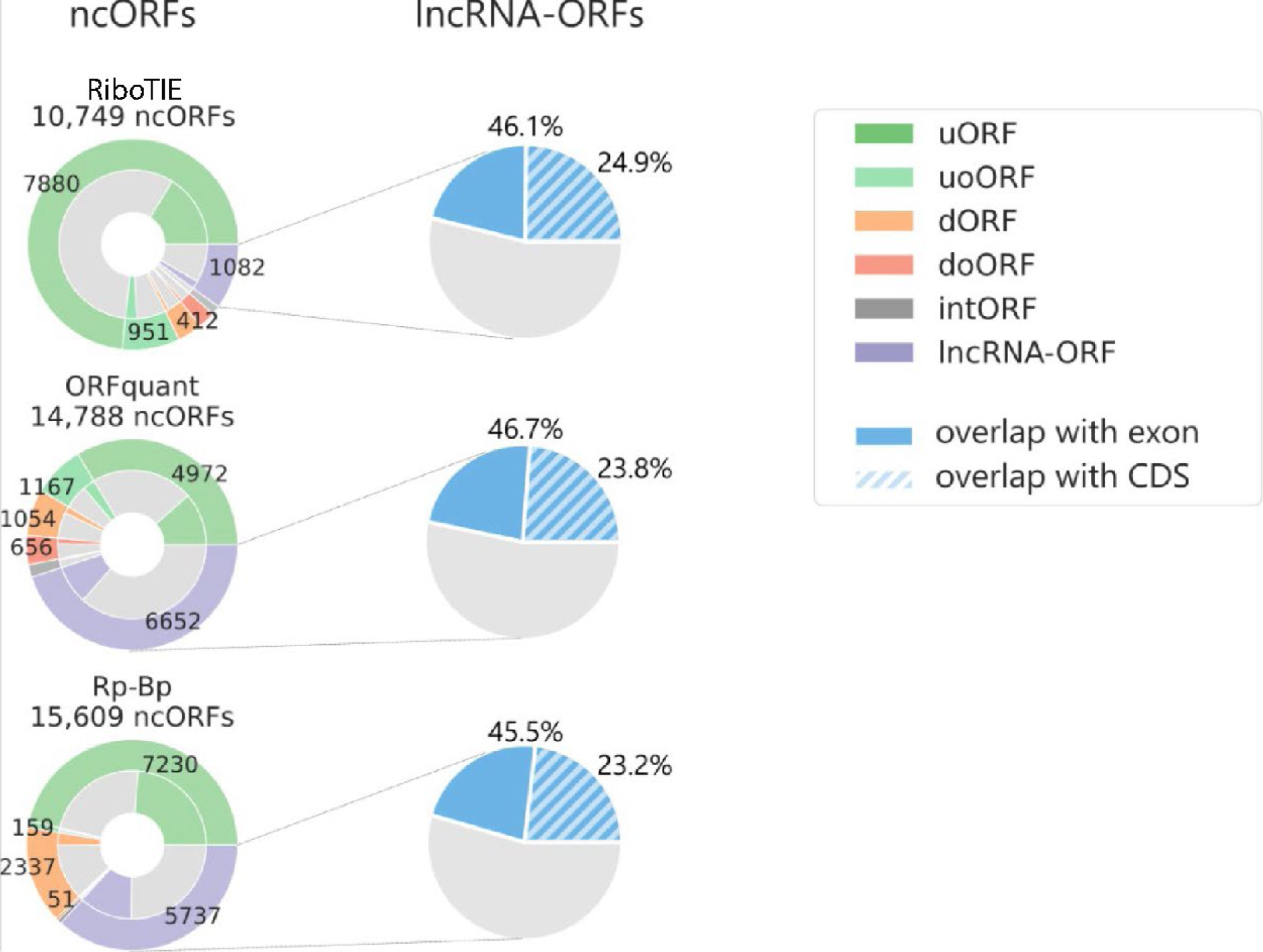
Characteristics of nominated lncRNA-ORFs by RiboTIE, ORFquant, and Rp-Bp on pancreatic progenitor cells. Overlap with protein coding exons and CDSs is evaluated using the TISs of nominated lncRNA-ORF.

**Extended Data Figure 6:** Example ncORFs with differential expression between medulloblastoma cell lines with high and low MYC expression. Given are the model outputs (y-axis) for positions of the transcript (x-axis) where the output of the model (RiboTIE: left; TIS Transformer: right) is larger than 0.04. The area of highly predicted ORFs is shown in orange (ncORF) and gray (annotated CDS). Ribo-Seq data, summed for all experiments in each group (low/high MYC), are displayed as orange bar plots (logarithmic scale, right y-axis).

**Extended Data Figure 7:**
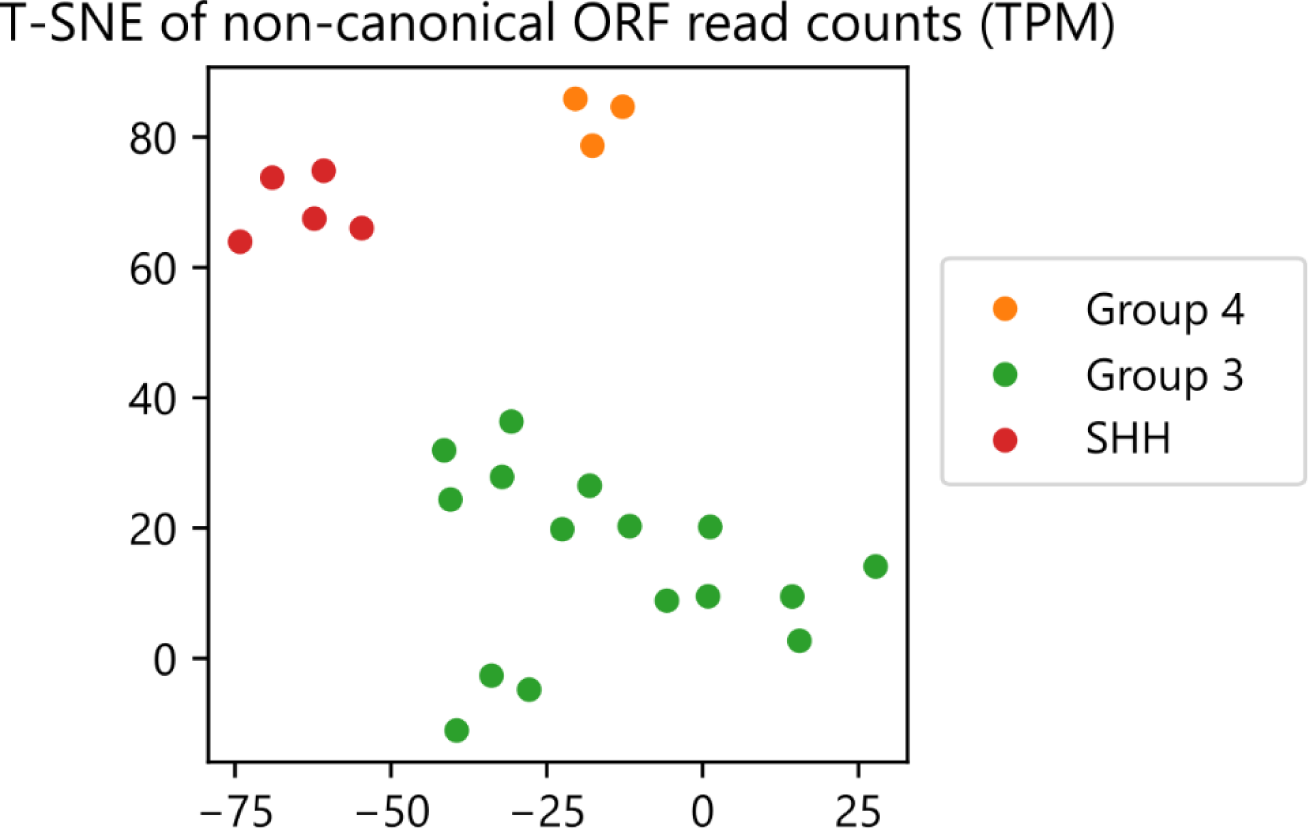
Clustering of Medulloblastoma cell line samples on non-canonical ORFs as called by RiboTIE. Clustering is performed on the normalized number of mapped reads (Transcripts Per Million (TPM)) using both PCA and T-SNE (Extended Data Table 4).

