## Supplementary Files for "Deep learning to decode sites of RNA translation in normal and cancerous tissues"

---

### 1. Supplementary Methods

#### 1.1. Ribosome profiling processing

Cutadapt and STAR are applied for trimming adapters and mapping reads to the genome and transcriptome. Read lengths between 20 and 40 nucleotides are retained. Reads mapping against tRNA/rRNA/sm(o)RNA are filtered out. Extended Data Table 1 lists the total number of reads within the dataset at various steps. The data were selected in order to have variation with respect to applied treatments and mapped number of reads.

```
# trim files and perform fastqc
cutadapt -j 20 -m 20 -a $adapter ${dataset}.fastq -o "out/temp/${dataset}_trimmed.fq" > "
    ↪ out/temp/${dataset}_trimmed_report.txt"

# remove rRNA/tRNA/smRNA/smoRNA
STAR --genomeLoad NoSharedMemory --seedSearchStartLmaxOverLread .5 --genomeDir '../../
    ↪ genome/STAR/excl_RNA' --readFilesIn $trimmed --outFilterMultimapNmax 1000 --
    ↪ outFilterMismatchNmax 2 --outFileNamePrefix out/temp/ --runThreadN 20 --
    ↪ outReadsUnmapped Fastx
mv out/temp/Unmapped.out.mate1 $cleaned

# align to genome, output mapping to transcriptome as well
STAR --runThreadN 20 --genomeDir '../../genome/STAR' --genomeLoad NoSharedMemory --
    ↪ readFilesIn $cleaned --outFileNamePrefix out/ --outSAMtype BAM SortedByCoordinate
    ↪ --quantMode TranscriptomeSAM --outSAMattributes MD NH --outFilterMultimapNmax 10
    ↪ --outMultimapperOrder Random --outFilterMismatchNmax 2 --
    ↪ seedSearchStartLmaxOverLread 0.5 --alignEndsType EndToEnd --outWigType bedGraph
```

#### 1.2. RiboTIE

##### 1.2.1. Data processing

Data loading for RiboTIE is achieved by storing data in the hierarchical data format version 5 (*hdf5*). Using Python, the ribosome reads mapped to the transcriptome are stored by transcript. The generated *bam* files are parsed using Python and data is stored to the *hdf5* format. Data is aggregated by the total number of reads aligned by their 5' position for every read length and transcript position. Transcript matrices are loaded from the *hdf5* files by a PyTorch data loader object and used as inputs to the model.

##### 1.2.2. Input embedding strategies

The transformer architecture takes full transcript regions as input and provides a prediction along each position of the input range. No sequence information is processed. No ORFs are identified as a pre-processing step. Transformer networks use vector representations of mapped reads at each nucleotide position as input tokens. As part of this research, different approaches were explored to create input vector representations from the mapped ribosome profiling data. For all instances, read counts are normalized for each transcript. This ensures the numerical stability of the inputs. Supplementary Figures 1 and Extended Data Figure 2 illustrate both strategies evaluated as part of this paper.

##### 1.2.3. Model architecture

The architectural framework is identical to that of TIS transformer[21], a transformer model used for predicting translation initiation sites using transcript sequence information. The transformer structure features multiple layers with multiple attention heads per layer. These are identical in structure but feature unique trainable model parameters. The outputs of the transformer module are sent to a set of fully connected layers to obtain a binary output at each input position. Notwithstanding the size of the dataset and overall high computational requirements of transformer architectures, model optimization **from scratch** is possible on a single RTX 3090 and converges after ca. 7 hours due to the relative shallowness of the transformer architecture as compared to many language-learning transformers.

##### 1.2.4. Attention

Custom attention strategies can be performed by the attention heads independent of the number of weights utilized to calculate the **Q**, **K**, **V** matrices. In this model, full attention is calculated through the Fast Attention Via Positive Orthogonal Random Features (FAVOR+) algorithm [22]. These allow full attention, where all inputs along the transcript are included by the attention head. In contrast, local attention restricts the attention matrix to only neighboring positions. Local attention is implemented by

**Algorithm 1** RiboTIE network architecture. Given are the different layers, their respective dimensions as defined by their hyperparameter names, the dimensions for RiboTIE (Table 5), and the resulting total weights. The bias term applied in each node is included and marked with italics.

- **RiboTIE** | **211,964**
  - **Ribosome Read Count** | **21,546**
    - **Linear** | $1 \times \text{dim}$ | $1 \times 42 + \mathit{42}$ | **84**
    - **Linear** | $\text{dim} \times \text{dim} * 6$ | $42 \times 252 + \mathit{252}$ | **10,836**
    - **Linear** | $\text{dim} * 6 \times \text{dim}$ | $252 \times 42 + \mathit{42}$ | **10,626**
  - **Ribosome Read Count Embedding** | $1 \times \text{dim}$ | $1 \times 42$ | **42**
  - **Ribosome Read Length Embedding** | read lengths $\times$ dim | $21 \times 42$ | **882**
  - **Positional Embedding** | fixed positional embeddings | **0**
  - **Performer** | **185,712**
    - **Layer** ($\times$ depth | 6) | **30,952**
      - **Layer norm** | $\text{dim} \times 2$ | $42 \times 2 + \mathit{2}$ | **86**
      - **Attention head** ($\times$ n_head | 6) | **2,064**
        - $\mathbf{W_Q}$ | dim $\times$ dim_head | $42 \times 16 + \mathit{16}$ | **688**
        - $\mathbf{W_K}$ | dim $\times$ dim_head | $42 \times 16 + \mathit{16}$ | **688**
        - $\mathbf{W_V}$ | dim $\times$ dim_head | $42 \times 16 + \mathit{16}$ | **688**
      - $\mathbf{W_o}$ | dim_head $*$ n_head $\times$ dim | $96 \times 42 + \mathit{42}$ | **4,074**
      - **Layer norm** | $\text{dim} \times 2$ | $42 \times 2 + \mathit{2}$ | **86**
      - **Linear** | $\text{dim} \times \text{dim} * 4$ | $42 \times 168 + \mathit{168}$ | **7,224**
      - **Linear** | $\text{dim} * 4 \times \text{dim}$ | $168 \times 42 + \mathit{42}$ | **7,098**
  - **Linear** | $\text{dim} \times \text{dim} * 2$ | $42 \times 84 + \mathit{84}$ | **3,612**
  - **Linear** | $\text{dim} \times 2$ | $84 \times 2 + \mathit{2}$ | **170**

dividing the attention matrix in smaller blocks on which full attention is calculated. Three blocks around the evaluated input are calculated. These local attention heads do not apply the FAVOR+ algorithm and use rotary positional embeddings [23]. The block size of the local attention heads is referred to under the 'attention scheme' columns of Supplementary Table 5.

#### 1.3. Training and Evaluation

This study explores the use of transformer models to detect translated open reading frames using ribosome profiling data. This is achieved by detecting translated initiation sites, constituting a binary-classification task. Model evaluations follow a standard deep learning set-up featuring a training, validation and test set. Data is grouped according to chromosomes to prevent identical profiles of ribosome reads, possible due to the existence of transcript isoforms, being separated between the training, validation or test set. The data used for the hyperparameter and input strategy selection are chromosomes 3, 4, 5, 6, 8, 9, 10, 11, 12, 13, 15, 16, 17, 18, 20, 21, 22, X, and Y for the training set, chromsomes 2 and 14 for the validation set, and chromosomes 1, 7, 13, and 19 for the test set. The data used for the pre-training strategy selection and benchmarking with previous tools feature two sets (folds) in order to cover the full transcriptome (within the test set). The first fold has chromosomes 3, 5, 7, 11, 13, 15, 19, 21, and X for the training set, chromosomes 1, 9, and 17 for the validation set, and chromosomes 2, 4, 6, 8, 10, 12, 14, 16, 18, 20, 22, and Y for the test set. The second fold has chromosomes 2, 6, 8, 10, 14, 16, 18, 22, and Y for the training set, chromosomes 4, 12, and 20 for the validation set, and chromosomes 1, 3, 5, 7, 9, 11, 13, 15, 17, 19, 21, and X for the test set. Binary cross-entropy is applied as the loss function. For all set-ups described in this paper, a learning rate of $10^{-3}$ is applied. The loss on the validation set indicates the optimal point of the model fit and is used for model selection (i.e. early stopping) to prevent overfitting on the training set. All reported performances are obtained on the test set.

##### 1.3.1. Hyperparameter selection

Hyperparameter optimization is performed to identify the optimal model architecture for this learning problem. The benchmark dataset featuring the highest number of mapped reads (SRR2733100) was selected to perform the hyperparameter selection on. No individual hyperparameters were observed to be more effective than others in improving performances. However, a correlation exists between the total number of model parameters and model performance. Eight unique architectures have been evaluated featuring varying configurations for the size of the hidden dimension, number of layers, number of attention heads per layer and the dimension of the attention head itself (Supplementary Table 5). Supplementary Figure 5 shows the validation loss at different epochs of the various architectures.

##### 1.3.2. Input token strategy

The input token strategies follows the same data allocations as the hyperparameter selection. To evaluate strategy A, reads were mapped by their 5' reads without offsets, and with offsets calculated by Plastid and RiboWaltz. Plastid and RiboWaltz have been selected for A/P-site offset calling as they are both popular methods specifically created for this task. Package versions are those listed as 'Last update' in Table 1. Scripts are executed following our custom folder structure.

**Plastid**

```
reformat_transcripts --annotation_files genome/Homo_sapiens.GRCh38.107.gff3 --
    ↪ annotation_format GFF3 --output_format GTF2 genome/Homo_sapiens.GRCh38.107.gtf2
metagene generate genome/plastid/ --landmark cds_start --annotation_files genome/
    ↪ Homo_sapiens.GRCh38.107.gtf2
```

```
psite genome/plastid/_rois.txt ribo/${dataset}/out/plastid/ --min_length 20 --max_length
    ↪ 41 --require_upstream --count_files ribo/${dataset}/out/genome/${dataset}_aligned.
    ↪ bam
```

**RiboWaltz**

```
library(riboWaltz)

metadata <- read.table('ribo/metadata.txt', header = FALSE, sep = "", dec = ".")
annotation_db <- create_annotation('genome/Homo_sapiens.GRCh38.107.gtf')
for (i in metadata$V1){
    reads_list <- bamtolist(bamfolder=sprintf("ribo/%s/out/", i), annotation=annotation_db
        ↪ )
    filtered_list <- length_filter(data=reads_list, length_filter_mode="custom", length_
        ↪ range=20:40)
    psite_offset <- psite(filtered_list)
    dir.create(sprintf("ribo/%s/out/ribowaltz", i))
    write.table(psite_offset, sprintf("ribo/%s/out/ribowaltz/riboWaltz_offsets.csv", i),
        ↪ sep="\t")
}
```

#### 1.3.3. Fine-tuning

Fine-tuning a trained model has two important advantages as compared to training a model from scratch–faster convergence of the validation loss, substantially reducing optimization times, and improved performances. Model pre-training follows the same data groupings to ensure models are exposed to the same transcripts during training at any stage. Both self-supervised and supervised pre-training have been evaluated using eight independent datasets (Extended Data Table 1). The supervised training objective proved to be the most effective (Supplementary Table A2). When applying RiboTIE for mapping translated open reading frames on the transcriptome, multiple models are trained that cover different parts of the transcriptome during training and model selection (i.e. training and validation set).

#### 1.3.4. Benchmark

Code snippets used to run various tools. ORFquant was run without use of any flags except those selecting the input and output files and is thus not listed.

**PRICE**

```
gedi -e IndexGenome -s genome/Homo_sapiens.GRCh38.dna.primary_assembly.fa -a genome/
    ↪ Homo_sapiens.GRCh38.107.gtf -f genome/price -nobowtie -nostar -nokallisto
gedi -e Price -reads ribo/${dataset}/out/genome/${dataset}_aligned.bam -genomic
    ↪ Homo_sapiens.GRCh38.107 -prefix ribo/${dataset}/out/price/ -progress -plot
```

**Rp-Bp**

```
prepare-rpbp-genome ../scripts/benchmark/rpbp_full.yml --star-options "--
    ↪ genomeSAindexNbases␣10" --mem 10G --num-cpus 4 --logging-level INFO --log-file
    ↪ genome/rpbp/rpbp-genome.log --write-unfiltered
run-all-rpbp-instances ribo/${dataset}/out/rpbp/rpbp.yml --num-cpus 30 --logging-level
    ↪ INFO --mem 50G
```

**Ribo-TISH**

```
ribotish quality -b ribo/${dataset}/out/genome/${dataset}_aligned.bam -g genome/
    ↪ Homo_sapiens.GRCh38.107.gtf -f ribo/${dataset}/out/ribotish/quality.pdf -r ribo/${
    ↪ dataset}/out/ribotish/offset.txt -o ribo/${dataset}/out/ribotish/quality.txt -l
    ↪ 20,41
ribotish predict -b ribo/${dataset}/out/genome/${dataset}_aligned.bam -g genome/
    ↪ Homo_sapiens.GRCh38.107.gtf -f genome/Homo_sapiens.GRCh38.dna.primary_assembly.fa
    ↪ -o ribo/${dataset}/out/ribotish/orfs.txt --ribopara ribo/${dataset}/out/ribotish/
    ↪ offset.txt
```

**ribotricer**

```
ribotricer prepare-orfs --gtf genome/Homo_sapiens.GRCh38.107.gtf --fasta genome/
    ↪ Homo_sapiens.GRCh38.dna.primary_assembly.fa --prefix genome/ribotricer/ribo
ribotricer detect-orfs --bam ribo/${dataset}/out/genome/${dataset}_aligned.bam --
    ↪ ribotricer_index genome/ribotricer/ribo_candidate_orfs.tsv --prefix ribo/${dataset
    ↪ }/out/ribotricer/
```

### 2. Supplementary Tables

Supplementary Table 1: **Various tools designed to detect expressed coding sequences using ribosome profiling data.** Methods with * require RNA-seq data. The list is non-exhaustive.

| **Method** | **Author** | **Calling** **A/P-site** | **ORF** | **Year** | **Last update** | **Language** |
|---|---|---|---|---|---|---|
| Ribotricer | [24] | Yes | Yes | 2019 | 1.3.3 (2023) | Python |
| Ribodeblur | [25] | Yes | No | 2018 | 2018 | Python |
| RiboWaltz | [26] | Yes | No | 2018 | 1.2.0 (2021) | R |
| RibORF | [27] | Yes | Yes | 2018 | 2.0 (2022) | Perl |
| RiboCode | [28] | Yes | Yes | 2018 | 1.2.15 (2022) | Python |
| RiboWave | [29] | Yes | Yes | 2018 | 2018 | Python |
| Scikit-ribo* | [30] | No | Yes | 2018 | 2018 | Python |
| Rp-Bp | [31] | Yes | Yes | 2017 | 3.0.1 (2023) | Python |
| RiboTISH | [32] | Yes | Yes | 2017 | 0.2.7 (2021) | Python |
| Plastid | [33] | Yes | No | 2016 | 0.6.1 (2022) | Python |
| PRICE (Gedi) | [34] | No | Yes | 2016 | 1.0.5 (2022) | Python |
| SPECtre | [35] | No | Yes | 2016 | 1.0.0 (2018) | R/Python |
| riboHMM* | [36] | No | Yes | 2016 | 2016 | Python |
| RiboProfiling | [37] | Yes | Yes | 2016 | 1.28.0 (2022) | R |
| RiboTaper* | [38] | No | Yes | 2015 | 1.3 (2016) | R |
| ORF-RATER | [39] | No | Yes | 2015 | 2018 | Python2.7 |
| PROTEOFORMER | [40] | Plastid | Yes | 2015 | 2.0 (2022) | Python2.7/Perl |

Supplementary Table 2: **A/P-offsets determined by various tools evaluated during this study.** The listed tools are Plastid (P), RiboWaltz (W), RiboCode (C), and RiboTISH (T)

| | **SRR1802129** | | | | | **SRR2433794** | | | | | **SRR2732970** | | | | | **SRR2733100** | | | | |
|---|---|---|---|---|---|---|---|---|---|---|---|---|---|---|---|---|---|---|---|---|
| | % | **P** | **W** | **C** | **T** | % | **P** | **W** | **C** | **T** | % | **P** | **W** | **C** | **T** | % | **P** | **W** | **C** | **T** |
| **20** | 0.2 | 13 | 7 | - | - | 0.1 | 13 | 13 | - | - | 1.3 | 2 | 12 | - | - | 0.8 | 2 | 12 | - | - |
| **21** | 0.2 | 13 | 9 | 5 | 17 | 0.2 | 13 | 13 | - | - | 1.6 | 3 | 12 | - | 12 | 1.0 | 3 | 12 | 12 | 12 |
| **22** | 0.2 | 13 | 9 | - | - | 0.4 | 5 | 12 | - | - | 1.7 | 12 | 12 | - | - | 1.0 | 3 | 12 | - | - |
| **23** | 0.2 | 13 | 7 | - | 7 | 0.7 | 6 | 12 | - | 12 | 2.2 | 12 | 12 | - | - | 1.2 | 5 | 12 | - | - |
| **24** | 0.3 | 13 | 8 | - | 8 | 1.0 | 7 | 12 | - | - | 2.2 | 12 | 12 | - | 12 | 1.4 | 5 | 12 | - | 12 |
| **25** | 0.4 | 13 | 9 | - | 12 | 2.4 | 9 | 12 | 12 | 12 | 2.3 | 12 | 12 | - | - | 1.5 | 12 | 12 | - | - |
| **26** | 1.7 | 50 | 10 | - | - | 5.2 | 9 | 12 | - | 12 | 2.5 | 12 | 12 | - | - | 1.8 | 12 | 12 | - | - |
| **27** | 9.6 | 12 | 12 | - | 11 | 8.7 | 10 | 12 | - | 12 | 3.1 | 12 | 12 | 12 | 12 | 2.5 | 12 | 12 | 12 | 12 |
| **28** | 36.3 | 12 | 12 | 12 | 12 | 19.7 | 12 | 12 | - | 12 | 4.8 | 12 | 12 | 12 | 12 | 4.3 | 12 | 12 | - | 12 |
| **29** | 32.0 | 12 | 13 | - | 12 | 37.4 | 12 | 12 | 12 | 12 | 14.5 | 12 | 12 | 12 | 12 | 13.3 | 12 | 12 | 12 | 12 |
| **30** | 9.0 | 23 | 15 | - | 13 | 21.8 | 12 | 12 | 12 | 12 | 30.0 | 12 | 12 | 12 | 12 | 27.7 | 12 | 12 | 12 | 12 |
| **31** | 1.7 | 13 | 14 | - | - | 2.1 | 13 | 13 | - | - | 21.6 | 12 | 12 | 12 | 12 | 25.0 | 12 | 12 | - | 12 |
| **32** | 1.4 | 13 | 16 | - | - | 0.3 | 50 | 11 | - | - | 8.3 | 12 | 13 | 12 | 12 | 12.5 | 13 | 13 | - | - |
| **33** | 2.5 | 13 | 17 | - | - | 0.1 | 13 | 12 | - | 15 | 2.5 | 13 | 13 | - | - | 4.0 | 5 | 13 | - | - |
| **34** | 2.5 | 13 | 19 | - | - | 0.0 | 13 | 14 | - | - | 0.8 | 12 | 13 | - | - | 1.0 | 5 | 13 | - | - |
| **35** | 1.4 | 13 | 18 | - | - | 0.0 | 13 | 13 | - | - | 0.3 | 13 | 13 | - | - | 0.4 | 13 | 14 | - | - |
| **36** | 0.1 | 13 | 20 | - | - | 0.0 | 13 | 15 | - | - | 0.2 | 12 | 12 | - | - | 0.2 | 13 | 12 | - | - |
| **37** | 0.1 | 13 | 21 | - | - | 0.0 | 13 | 12 | - | - | 0.1 | 50 | 12 | - | - | 0.2 | 13 | 14 | - | - |
| **38** | 0.1 | 13 | 22 | - | - | 0.0 | 13 | 12 | - | - | 0.1 | 13 | 12 | - | - | 0.1 | 13 | 12 | - | - |
| **39** | 0.1 | 13 | 23 | - | - | 0.0 | 13 | 12 | - | - | 0.0 | 50 | 12 | - | - | 0.1 | 13 | 14 | - | - |
| **40** | 0.0 | 13 | 24 | - | - | 0.0 | 13 | 12 | - | - | 0.0 | 13 | 12 | - | - | 0.0 | 13 | 13 | - | - |

| | **SRR2954800** | | | | | **SRR8449577** | | | | | **SRR9113067** | | | | | **SRR11005875** | | | | |
|---|---|---|---|---|---|---|---|---|---|---|---|---|---|---|---|---|---|---|---|---|
| | % | **P** | **W** | **C** | **T** | % | **P** | **W** | **C** | **T** | % | **P** | **W** | **C** | **T** | % | **P** | **W** | **C** | **T** |
| **20** | 0.0 | 13 | 7 | - | 9 | 1.0 | 5 | 12 | - | - | 4.6 | 3 | 12 | - | 12 | 0.1 | 13 | 12 | - | 12 |
| **21** | 0.0 | 13 | 11 | - | 8 | 1.6 | 5 | 12 | - | 12 | 6.8 | 12 | 12 | 12 | 12 | 0.4 | 13 | 12 | - | - |
| **22** | 0.1 | 11 | 11 | - | - | 1.6 | 6 | 12 | 12 | 12 | 4.7 | 29 | 13 | - | - | 0.8 | 8 | 12 | - | - |
| **23** | 0.1 | 13 | 12 | - | - | 1.7 | 7 | 12 | - | - | 3.2 | 3 | 12 | - | 12 | 1.3 | 6 | 12 | - | 12 |
| **24** | 0.6 | 5 | 11 | - | 11 | 2.0 | 8 | 12 | - | - | 3.1 | 3 | 12 | - | 12 | 1.7 | 7 | 12 | - | - |
| **25** | 1.4 | 6 | 12 | - | - | 2.9 | 9 | 12 | 12 | 12 | 3.5 | 12 | 12 | - | 12 | 2.1 | 8 | 12 | - | - |
| **26** | 2.1 | 7 | 11 | - | - | 6.3 | 11 | 12 | - | - | 4.3 | 3 | 12 | - | 12 | 3.1 | 9 | 12 | - | 12 |
| **27** | 3.3 | 11 | 11 | 8 | 11 | 15.7 | 12 | 12 | - | - | 5.4 | 12 | 12 | - | 12 | 5.3 | 11 | 12 | - | - |
| **28** | 4.7 | 11 | 11 | - | - | 43.2 | 12 | 12 | 12 | 12 | 8.8 | 12 | 12 | - | 12 | 23.5 | 12 | 12 | 12 | 12 |
| **29** | 10.5 | 11 | 11 | 11 | 11 | 19.2 | 12 | 12 | 12 | 12 | 17.4 | 12 | 12 | 12 | 12 | 45.7 | 12 | 12 | 12 | 12 |
| **30** | 26.9 | 11 | 12 | - | 12 | 2.5 | 6 | 12 | 12 | 12 | 19.0 | 12 | 12 | - | 12 | 13.7 | 12 | 12 | 12 | 12 |
| **31** | 21.0 | 12 | 12 | - | 12 | 0.9 | 13 | 12 | 12 | 12 | 11.5 | 13 | 13 | - | 12 | 1.5 | 50 | 12 | - | - |
| **32** | 9.8 | 12 | 12 | - | 12 | 0.5 | 13 | 12 | - | 12 | 4.9 | 10 | 13 | - | - | 0.3 | 13 | 12 | - | - |
| **33** | 5.9 | 11 | 12 | - | - | 0.4 | 13 | 12 | - | - | 2.0 | 13 | 11 | - | - | 0.2 | 13 | 12 | - | - |
| **34** | 4.5 | 12 | 12 | - | - | 0.2 | 13 | 13 | 12 | - | 0.5 | 13 | 14 | - | - | 0.1 | 13 | 12 | - | - |
| **35** | 3.4 | 13 | 12 | - | - | 0.1 | 13 | 12 | - | - | 0.2 | 13 | 12 | - | - | 0.0 | 13 | 12 | - | 12 |
| **36** | 2.0 | 50 | 12 | - | - | 0.1 | 13 | 14 | - | - | 0.1 | 13 | 11 | - | - | 0.0 | 13 | 10 | - | - |
| **37** | 1.2 | 13 | 11 | - | - | 0.0 | 13 | 11 | - | - | 0.1 | 13 | 11 | - | - | 0.0 | 13 | 18 | - | - |
| **38** | 1.0 | 13 | 13 | - | - | 0.0 | 13 | 12 | - | - | 0.1 | 13 | 12 | - | - | 0.0 | 13 | 12 | - | - |
| **39** | 1.1 | 13 | 11 | - | - | 0.0 | 13 | 12 | - | - | 0.0 | 13 | 12 | - | - | 0.0 | 13 | 12 | - | - |
| **40** | 0.3 | 13 | 13 | - | - | 0.0 | 13 | 12 | - | - | - | 13 | 12 | - | - | 0.0 | 13 | 12 | - | - |

Supplementary Table 3: **RiboTIE performances for different input token strategies and datasets**. Scores are calculated on the test set after selection of the model with the minimum validation loss (See Extended Data Figure 2). For each dataset and strategy, the cross-entropy loss ($\times 10^3$), area under the receiver operating characteristic curve (ROC), and area under the precision-recall curve (PR) are given. Results indicate the relevance of read length information for the prediction of translation initiation sites using ribosome profiling data, especially for datasets featuring a higher read depth (see Extended Data Table 1). All strategies are evaluated using the same model architecture and training/validation data (Architecture 4, see Supplementary Table 5, Supplementary Figure 5), Strategy A generates input tokens utilizing read count information for every position of the transcript. Strategy A includes mappings generated by taking the 5' position of every read, and offsetting reads based on read length utilizing two different tools (Plastid, RiboWaltz). Strategy B includes information on both the positions and read lengths of the mapped reads.

| | | **SRR1802129** | | | **SRR2433794** | | | **SRR2732970** | | | **SRR2733100** | | |
|---|---|---|---|---|---|---|---|---|---|---|---|---|---|
| | **Position** | **Loss** | **ROC** | **PR** | **Loss** | **ROC** | **PR** | **Loss** | **ROC** | **PR** | **Loss** | **ROC** | **PR** |
| **A** | 5' | 1.71 | 0.938 | 0.0211 | 1.48 | 0.965 | 0.0937 | 1.40 | 0.963 | 0.161 | 1.38 | 0.965 | 0.161 |
| **A** | Plastid | 1.74 | 0.935 | 0.0144 | 1.49 | 0.964 | 0.0825 | 1.39 | 0.965 | 0.156 | 1.41 | 0.964 | 0.145 |
| **A** | RiboWaltz | 1.7 | 0.941 | 0.0205 | 1.47 | 0.966 | 0.0908 | 1.4 | 0.964 | 0.154 | 1.4 | 0.964 | 0.151 |
| **B** | 5' | **1.69** | 0.945 | 0.0217 | **1.44** | 0.968 | 0.104 | **1.31** | 0.969 | 0.211 | **1.3** | 0.97 | 0.217 |

| | | **SRR2954800** | | | **SRR8449577** | | | **SRR9113067** | | | **SRR11005875** | | |
|---|---|---|---|---|---|---|---|---|---|---|---|---|---|
| | **Position** | **Loss** | **ROC** | **PR** | **Loss** | **ROC** | **PR** | **Loss** | **ROC** | **PR** | **Loss** | **ROC** | **PR** |
| **A** | 5' | 1.69 | 0.935 | 0.0394 | 1.55 | 0.955 | 0.0721 | 1.7 | 0.943 | 0.0178 | 1.48 | 0.967 | 0.0727 |
| **A** | Plastid | 1.72 | 0.931 | 0.0309 | 1.57 | 0.954 | 0.0579 | 1.7 | 0.942 | 0.0161 | 1.5 | 0.966 | 0.0721 |
| **A** | RiboWaltz | 1.71 | 0.932 | 0.0324 | 1.56 | 0.955 | 0.064 | 1.7 | 0.943 | 0.0171 | 1.48 | 0.967 | 0.0721 |
| **B** | 5' | **1.69** | 0.937 | 0.04 | **1.54** | 0.956 | 0.0751 | **1.7** | 0.943 | 0.0189 | **1.47** | 0.967 | 0.0804 |

Supplementary Table 4: **RiboTIE performances for different model optimization strategies and datasets**. Scores are calculated on the test set after selection of the model with the minimum validation loss (See Supplementary Figure 7, 8). For each dataset and strategy, the cross-entropy loss ($\times 10^3$), area under the receiver operating characteristic curve (ROC), and area under the precision-recall curve (PR) are given. Results show the gain by having a model trained on a large variety of ribosome-profiling datasets using a supervised learning objective (see Extended Data Table 1). All settings are evaluated using the same model architecture (Architecture 4, see Supplementary Table 5) and input token strategy (Strategy B, see Supplementary Table 3. The data is split in two folds (F1 and F2), with different parts of the transcriptome covered as training/validation/test data in each fold.

| | | **SRR1802129** | | | **SRR2433794** | | | **SRR2732970** | | | **SRR2733100** | | |
|---|---|---|---|---|---|---|---|---|---|---|---|---|---|
| | **Pre-train** | **Loss** | **ROC** | **PR** | **Loss** | **ROC** | **PR** | **Loss** | **ROC** | **PR** | **Loss** | **ROC** | **PR** |
| | - | 1.58 | 0.944 | 0.017 | 1.36 | 0.968 | 0.097 | 1.24 | 0.969 | 0.193 | 1.22 | 0.969 | 0.203 |
| **F1** | Supervised | **1.56** | 0.948 | 0.024 | **1.32** | 0.970 | 0.120 | **1.18** | 0.972 | 0.239 | **1.18** | 0.972 | 0.240 |
| | Self-Supervised | 1.57 | 0.946 | 0.020 | 1.36 | 0.966 | 0.110 | 1.20 | 0.970 | 0.227 | 1.20 | 0.969 | 0.226 |
| | - | 1.66 | 0.944 | 0.015 | 1.47 | 0.962 | 0.084 | 1.29 | 0.968 | 0.193 | 1.29 | 0.969 | 0.195 |
| **F2** | Supervised | **1.63** | 0.948 | 0.024 | **1.42** | 0.967 | 0.104 | **1.25** | 0.972 | 0.227 | **1.24** | 0.972 | 0.229 |
| | Self-Supervised | 1.65 | 0.945 | 0.021 | 1.44 | 0.965 | 0.098 | 1.26 | 0.969 | 0.214 | 1.28 | 0.969 | 0.214 |

| | | **SRR2954800** | | | **SRR8449577** | | | **SRR9113067** | | | **SRR11005875** | | |
|---|---|---|---|---|---|---|---|---|---|---|---|---|---|
| | **Position** | **Loss** | **ROC** | **PR** | **Loss** | **ROC** | **PR** | **Loss** | **ROC** | **PR** | **Loss** | **ROC** | **PR** |
| | - | 1.60 | 0.935 | 0.033 | 1.43 | 0.959 | 0.079 | 1.59 | 0.946 | 0.013 | 1.37 | 0.969 | 0.075 |
| **F1** | Supervised | **1.57** | 0.939 | 0.045 | **1.40** | 0.962 | 0.096 | **1.55** | 0.951 | 0.024 | **1.34** | 0.972 | 0.094 |
| | Self-Supervised | 1.59 | 0.936 | 0.036 | 1.42 | 0.959 | 0.087 | 1.58 | 0.946 | 0.019 | 1.37 | 0.968 | 0.077 |
| | - | 1.70 | 0.932 | 0.025 | 1.52 | 0.956 | 0.071 | 1.68 | 0.941 | 0.013 | 1.45 | 0.966 | 0.076 |
| **F2** | Supervised | **1.65** | 0.939 | 0.044 | **1.47** | 0.960 | 0.092 | **1.62** | 0.949 | 0.027 | **1.42** | 0.970 | 0.090 |
| | Self-Supervised | 1.68 | 0.933 | 0.033 | 1.50 | 0.958 | 0.085 | 1.67 | 0.944 | 0.015 | 1.45 | 0.967 | 0.079 |

Supplementary Table 5: **Eight model architectures used for hyperparameter tuning selection.** For each set-up, a model is trained to detect translation initiation sites using ribosome profiling data. Hyperparameter selection is based on the minimum loss on the validation set. Hyperparameter tuning is performed on SRR2733100, featuring the highest read depth of all evaluated datasets. Supplementary Figure 5 displays the validation loss curves for each of the listed architectures.

| | **Hidden** | | **Attention head** | | **Attention scheme** | | | |
|---|---|---|---|---|---|---|---|---|
| **ID** | **state dim.** | **Depth** | **Heads** | **Head dim.** | **Local** | **Full** | **Model parameters** | **Val. loss ($\times 10^{-3}$)** |
| 1 | 24 | 5 | 6 | 12 | 4 | 2 | 81K | 1.099 |
| 2 | 30 | 6 | 6 | 16 | 4 | 2 | 129K | 1.105 |
| 3 | 30 | 8 | 6 | 24 | 4 | 2 | 215K | 1.108 |
| **4** | **42** | **6** | **6** | **16** | **4** | **2** | **211K** | **1.095** |
| 5 | 42 | 8 | 6 | 24 | 4 | 2 | 339K | 1.099 |
| 6 | 48 | 6 | 8 | 16 | 5 | 3 | 297K | 1.096 |
| 7 | 48 | 8 | 8 | 24 | 5 | 3 | 484K | 1.110 |
| 8 | 50 | 10 | 10 | 24 | 6 | 4 | 525K | 1.102 |

Supplementary Table 6: **Comparative performances of RiboTIE based on different subsets of the data**. The model predictions to detect translation initiation sites for each position on the trancriptome can be subsetted as a post-processing step. Using Ensembl translation initiation sites to derive a positive set, the area under the receiver operating characteristic curve (ROC) and area under the precision-recall curve (PR) are calculated. Given is the performance for all positions (total of ∼430M), with no conditions for what a valid ORF constitutes, positions that result in an ORF with a valid stop codon on the transcript (stop codon), an ORF length larger than 30 nucleotides (ORF length), and an ATG start codon (ATG start). Additionally, a subset has been selected using a minimum of 20 mapped reads (# Reads) on the transcript as a requirement. The performance and percentage of the total samples when using a combination of all listed conditions (Combined) is listed in the last set of columns. Note that the predictions of RiboTIE are those of the models pre-trained using a supervised learning strategy (see Supplementary Table 2), where the predictions of both models/folds are simply merged to cover the full transcriptome.

| | **-** | | **Stop codon** | | **ORF length** | | **# Reads** | | **ATG start** | | **Combined** | | |
|---|---|---|---|---|---|---|---|---|---|---|---|---|---|
| **dataset** | **ROC** | **PR** | **ROC** | **PR** | **ROC** | **PR** | **ROC** | **PR** | **ROC** | **PR** | **%** | **ROC** | **PR** |
| SRR1802129 | 0.945 | 0.020 | 0.946 | 0.020 | 0.943 | 0.021 | 0.981 | 0.041 | 0.952 | 0.318 | 0.6 | 0.983 | 0.502 |
| SRR2433794 | 0.965 | 0.101 | 0.966 | 0.103 | 0.964 | 0.103 | 0.983 | 0.137 | 0.966 | 0.419 | 1.0 | 0.981 | 0.516 |
| SRR2732970 | 0.969 | 0.220 | 0.970 | 0.222 | 0.968 | 0.223 | 0.986 | 0.285 | 0.958 | 0.399 | 1.1 | 0.972 | 0.487 |
| SRR2733100 | 0.969 | 0.215 | 0.970 | 0.217 | 0.968 | 0.218 | 0.986 | 0.279 | 0.957 | 0.399 | 1.1 | 0.971 | 0.488 |
| SRR2954800 | 0.935 | 0.034 | 0.935 | 0.034 | 0.932 | 0.036 | 0.972 | 0.068 | 0.942 | 0.266 | 0.6 | 0.974 | 0.417 |
| SRR8449577 | 0.958 | 0.083 | 0.959 | 0.084 | 0.957 | 0.085 | 0.983 | 0.128 | 0.961 | 0.382 | 0.8 | 0.982 | 0.514 |
| SRR9113067 | 0.944 | 0.016 | 0.945 | 0.016 | 0.942 | 0.017 | 0.971 | 0.024 | 0.950 | 0.289 | 0.9 | 0.974 | 0.399 |
| SRR11005875 | 0.967 | 0.077 | 0.968 | 0.078 | 0.966 | 0.080 | 0.985 | 0.106 | 0.969 | 0.432 | 1.0 | 0.984 | 0.534 |

### 3. Supplementary Figures

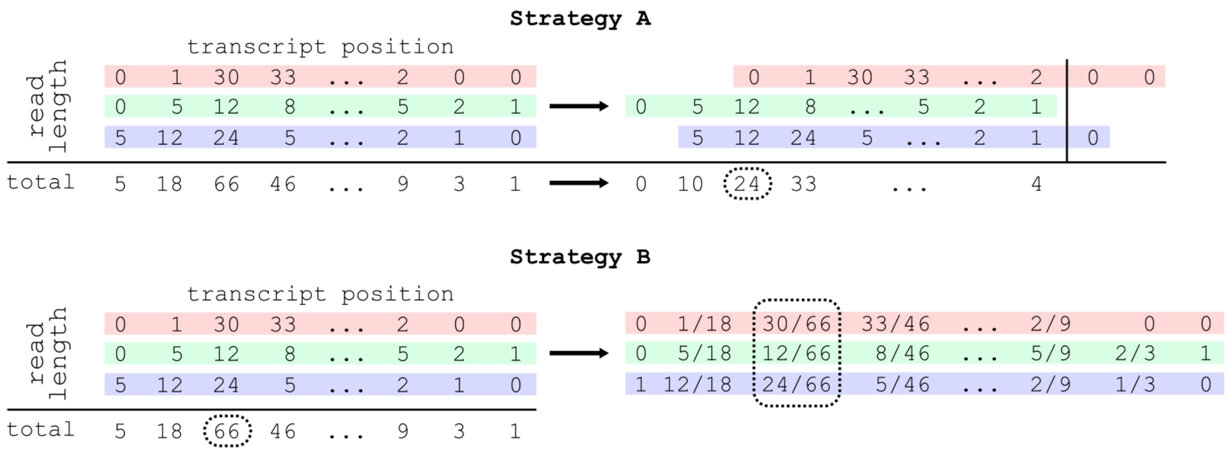


Supplementary Figure 1: **Illustration of the data applied for calculating the input vector representation.** For a given matrix containing reads mapped according to their 5'-end by transcript position and read length. Strategy A: reads are offset according to a fixed value for each read length. The total read count is applied for further processing. Strategy B: both the total read count and the fractional abundance of each read length is used to obtain an input vector representation. Input vector representations are calculated for each position (e.g. dotted square encapsulates data used for a single position). Note that in contrast to the illustration, data is generally sparse and ribosome profiling data is applied for 21 read lengths ($[20, 40]$).

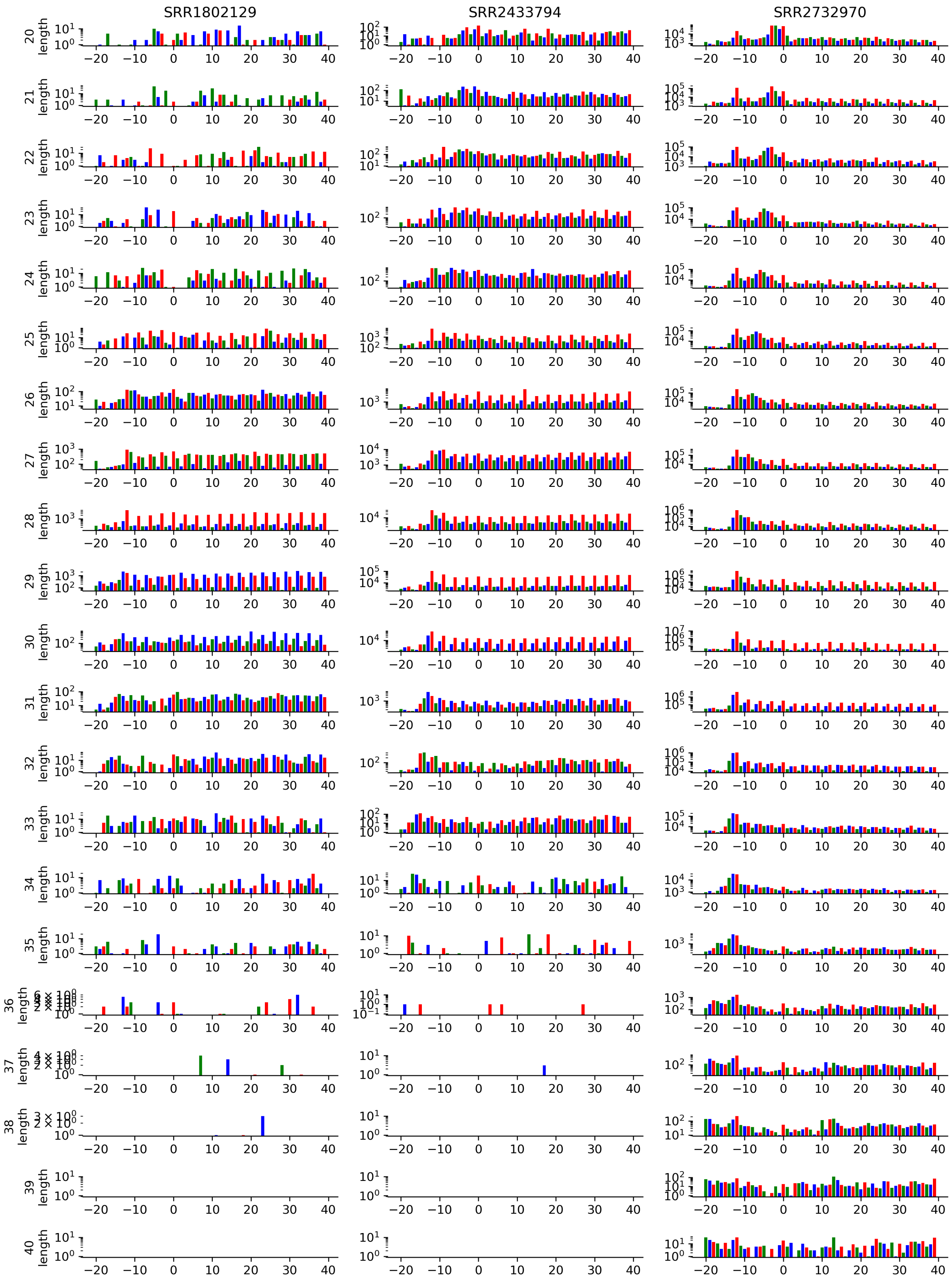


Supplementary Figure 2: **Counts of reads mapped by their 5' positions along translation initiation start sites.** The figure showcases unique patterns of read alignments per read length and experiment. Read counts are taken by only evaluating translation initiation sites of coding sequences within the consensus coding sequence (CCDS) library. A window of 20 nucleotides upstream and 40 nucleotides downstream is taken. A logarithmic scale and alternating color scheme is used to highlight the patterns emerging from the triplet periodicity along the translation initiation site and coding sequence. Included are experiments SRR1802129, SRR2433794, and SRR2732970. Accompanied by Supplementary Figure 3 and 4.

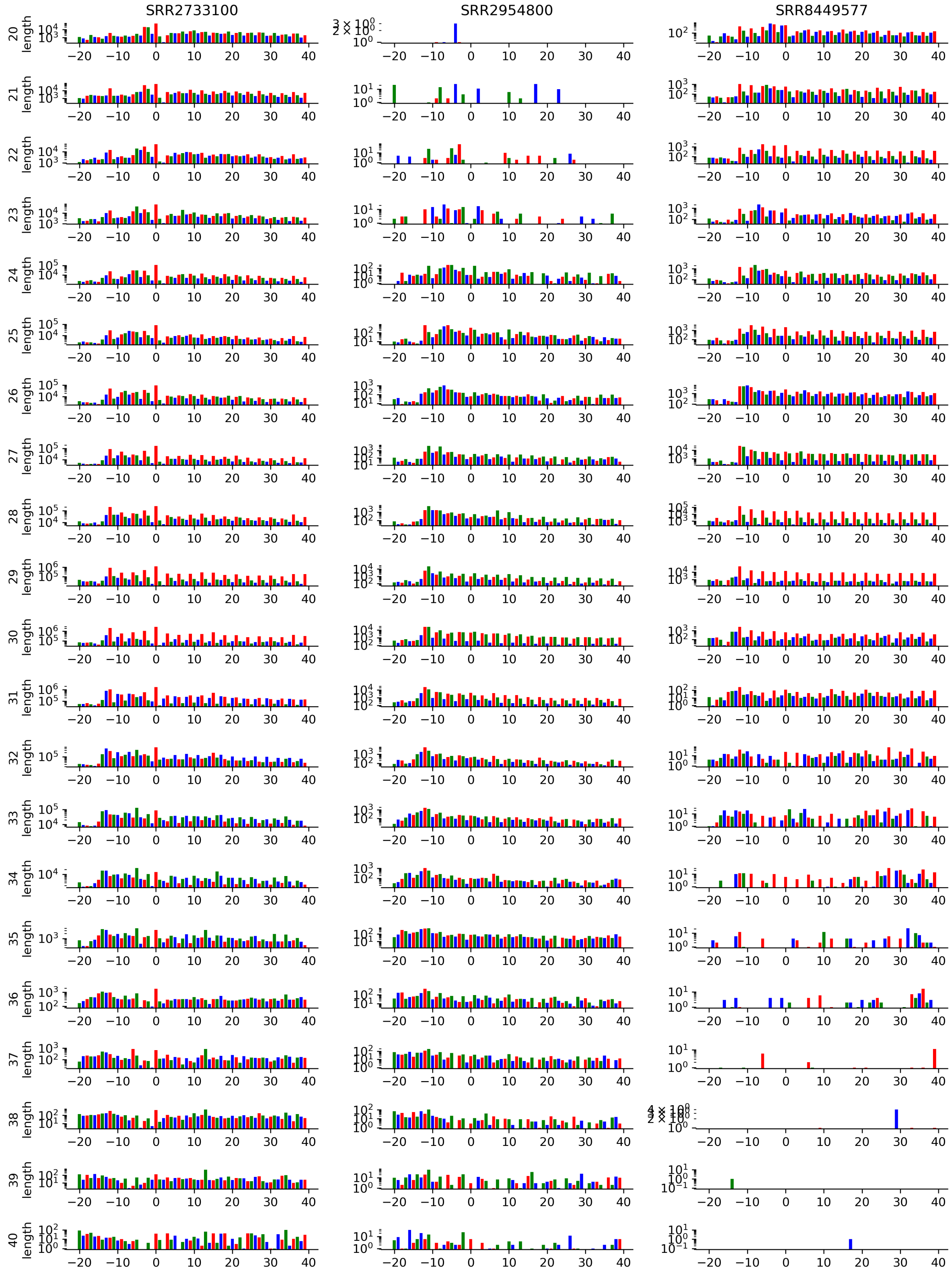


Supplementary Figure 3: **Counts of reads mapped by their 5' positions along translation initiation start sites.** The figure showcases unique patterns of read alignments per read length and experiment. Read counts are taken by only evaluating translation initiation sites of coding sequences within the consensus coding sequence (CCDS) library. A window of 20 nucleotides upstream and 40 nucleotides downstream is taken. A logarithmic scale and alternating color scheme is used to highlight the patterns emerging from the triplet periodicity along the translation initiation site and coding sequence. Included are experiments SRR2733100, SRR2954800, and SRR8449577. Accompanied by Supplementary Figure 2 and 4.

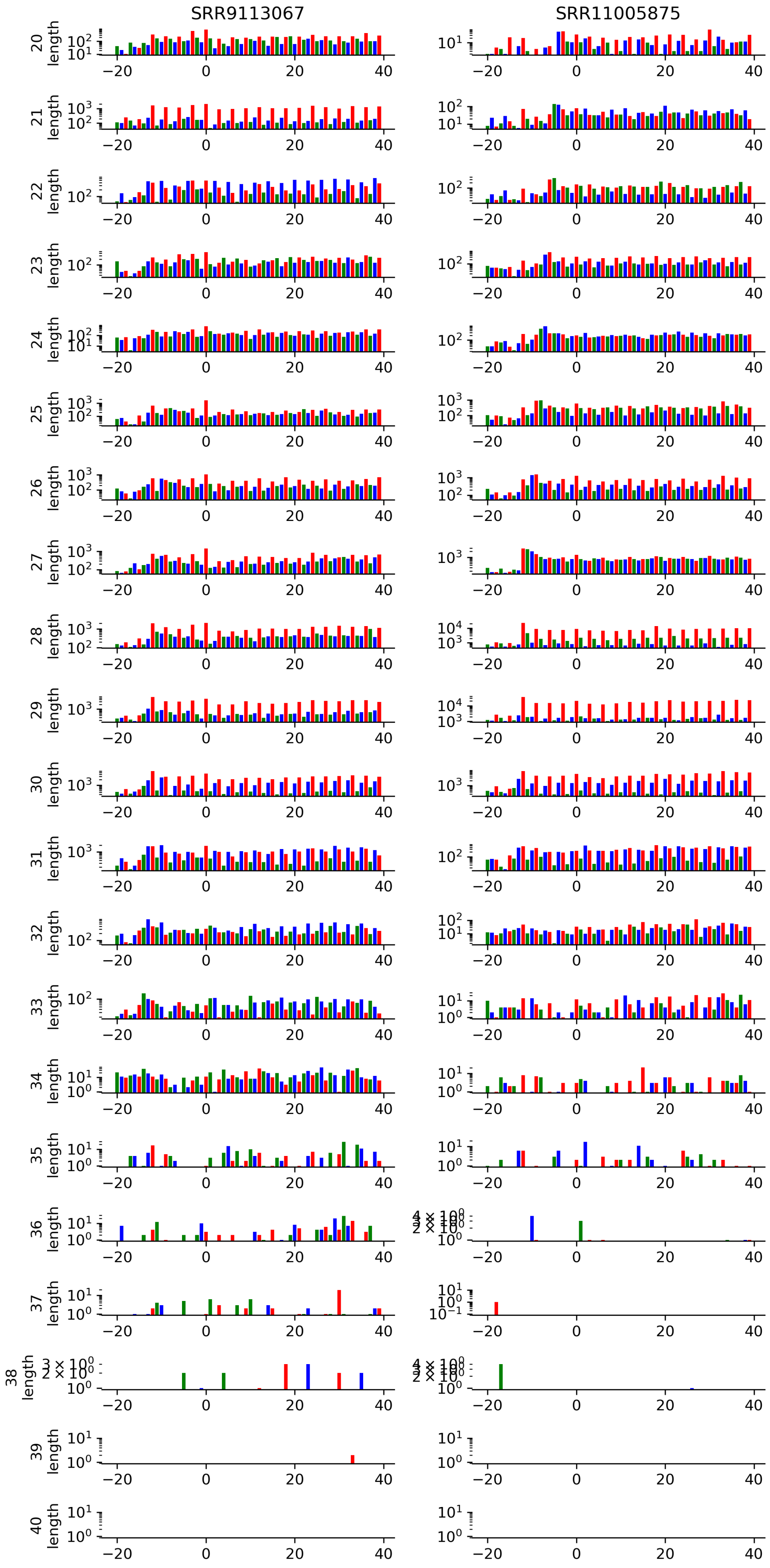


Supplementary Figure 4: **Counts of reads mapped by their 5' positions along translation initiation start sites.** The figure showcases unique patterns of read alignments per read length and experiment. Read counts are taken by only evaluating translation initiation sites of coding sequences within the consensus coding sequence (CCDS) library. A window of 20 nucleotides upstream and 40 nucleotides downstream is taken. A logarithmic scale and alternating color scheme is used to highlight the patterns emerging from the triplet periodicity along the translation initiation site and coding sequence. Included are experiments SRR9113067, SRR11005875. Accompanied by Supplementary Figure 2 and 3.

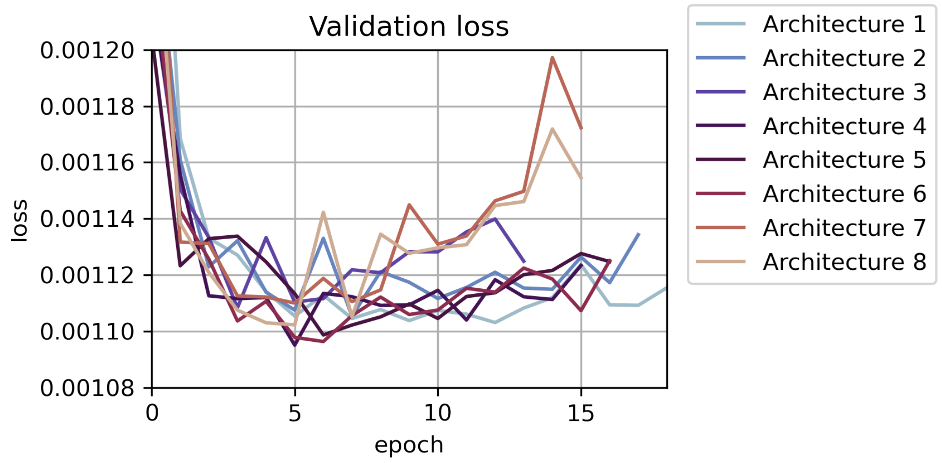


Supplementary Figure 5: **The loss curves of the model architectures trained for detecting TIS using ribosome profiling data.** The validation sets used are chromosomes 2 and 14. The hyperparameters for each model are given in Table 5. Architectures 1–2 converge slowly over several epochs without reaching a minimum within the evaluated time frame, indicating too few model weights. The higher number of model weights of architectures 7–8 result in clear overfitting from epoch 5 onward. Architecture 4 returns the lowest loss, and is selected for model benchmarking. While the minimum loss is similar for all architectures, the plot confirms our selection of a model architecture with a suitable number of parameters.

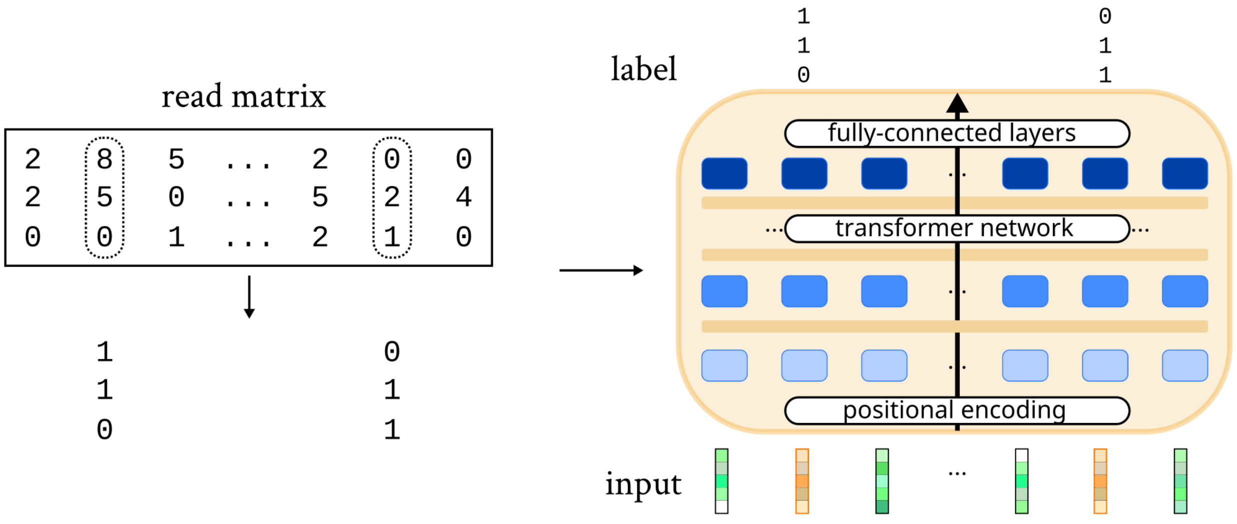


Supplementary Figure 6: **Self-supervised learning implementation for ribosome profiling data.** One of the pre-training approaches investigated in this paper. A model is trained to infer the presence of a mapped ribosome reads at a given position. The task constitutes a binary multi-label classification task. 15% of the input positions were randomly selected (dotted frame) and masked using a custom input embedding (orange input vectors). Positive labels are allocated to read lengths having more than one read mapped at a given position. Note that in contrast to the illustration, data is generally sparse and ribosome profiling data is applied for 21 read lengths ($[20, 40]$).

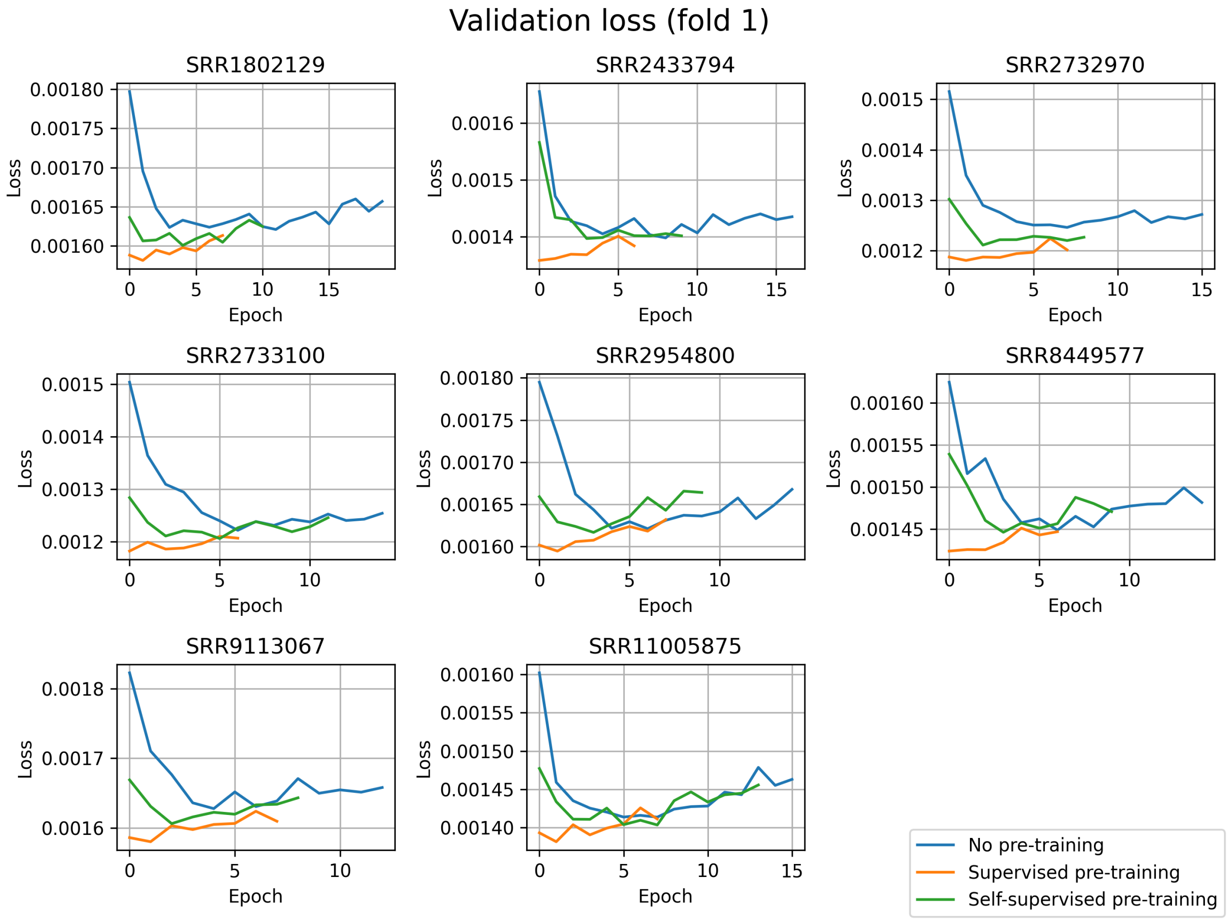


Supplementary Figure 7: **Validation cross-entropy loss of different training schemes of RiboTIE.** Eight datasets were evaluated following three approaches. This is achieved by training a model from scratch (no pre-training) or using a pre-trained model fit on a selection of eight separate datasets (see Extended Data Table 1). Pre-trained models include both those fit following a supervised learning objective (supervised pre-training) on identifying translation initiation sites, and those fit following a self-supervised learning objective, similar to those found in language processing (see Supplementary Figure 6). This figure shows the models trained on chromosomes 3, 5, 7, 11, 13, 15, 19, 21, and X with chromosomes 1, 9, and 17 used as validation set.

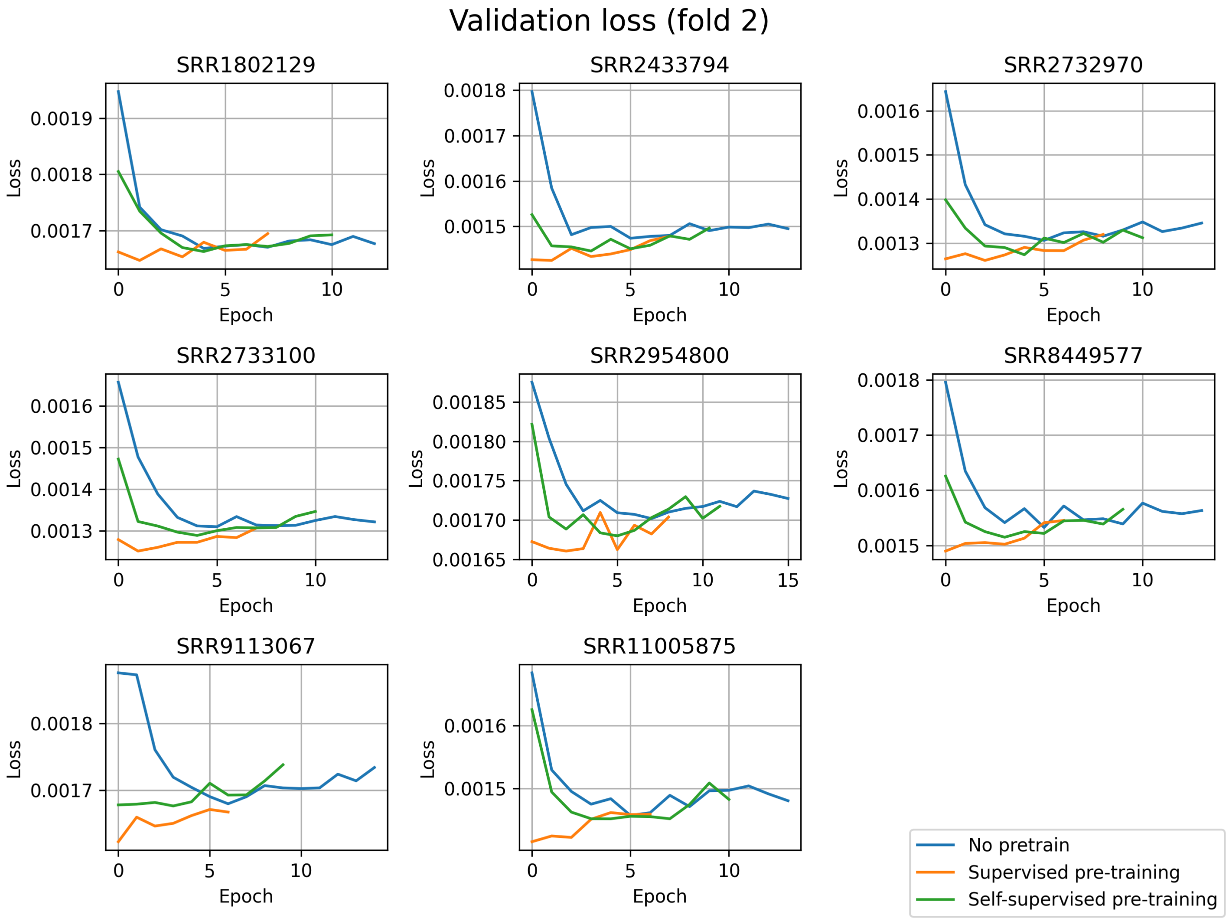


Supplementary Figure 8: **Validation cross-entropy loss of different training schemes of RiboTIE.** Eight datasets were evaluated following three approaches. This is achieved by training a model from scratch (no pre-training) or using a pre-trained model fit on a selection of eight separate datasets (see Extended Data Table 1). Pre-trained models include both those fit following a supervised learning objective (supervised pre-training) on identifying translation initiation sites, and those fit following a self-supervised learning objective, similar to those found in language processing (see Supplementary Figure 6). This figure shows the models trained on chromosomes 2, 6, 8, 10, 14, 16, 18, 22, and Y with chromosomes 4, 12, and 20 used as validation set.
